# AlphaFold-SFA: accelerated sampling of *cryptic pocket* opening, *protein-ligand* binding and *allostery* by AlphaFold, slow feature analysis and metadynamics

**DOI:** 10.1101/2023.11.21.568098

**Authors:** Shray Vats, Raitis Bobrovs, Pär Söderhjelm, Soumendranath Bhakat

## Abstract

Sampling *rare events* in proteins is crucial for comprehending complex phenomena like cryptic pocket opening, where transient structural changes expose new binding sites. Understanding these rare events also sheds light on protein-ligand binding and allosteric communications, where distant site interactions influence protein function. Traditional unbiased molecular dynamics simulations often fail to sample such rare events, as the free energy barrier between metastable states is large relative to the thermal energy. This renders these events inaccessible on the timescales typically simulated by standard molecular dynamics, limiting our understanding of these critical processes. In this paper, we proposed a novel unsupervised learning approach termed as *slow feature analysis* (SFA) which aims to extract slowly varying features from high-dimensional temporal data. SFA trained on small unbiased molecular dynamics simulations launched from AlphaFold generated conformational ensembles manages to capture rare events governing cryptic pocket opening, protein-ligand binding, and allosteric communications in a kinase. Metadynamics simulations using SFA as collective variables manage to sample ‘deep’ cryptic pocket opening within a few hundreds of nanoseconds which was beyond the reach of microsecond long unbiased molecular dynamics simulations. SFA augmented metadynamics also managed to capture accelerated ligand binding/unbinding and provided novel insights into allosteric communication in receptor-interacting protein kinase 2 (RIPK2) which dictates protein-protein interaction. Taken together, our results show how SFA acts as a dimensionality reduction tool which bridges the gap between AlphaFold, molecular dynamics simulation and metadynamics in context of capturing rare events in biomolecules, extending the scope of structure-based drug discovery in the era of AlphaFold.

## Introduction

The challenge of accurately predicting the 3D structure of proteins based solely on their amino acid sequences has been a longstanding puzzle in the realm of structural biology. Traditionally, scientists relied heavily on experimental techniques like X-ray crystallography and cryogenic electron microscopy (Cryo-EM) to decipher these protein structures^1^. While these methods remain crucial for examining intricate biomolecules, the landscape underwent a transformative shift in 2021. This change was marked by the introduction of AlphaFold^2^, an artificial intelligence (AI)-driven model that showcased its prowess in predicting protein 3D structures from their sequences. Building on this innovation, ColabFold^3^ was subsequently developed, optimizing AlphaFold to operate seamlessly on Google Colab, thereby democratizing AI-based protein structure prediction. However, it’s essential to highlight a shared limitation across AlphaFold, X-ray crystallography, and Cryo-EM: while they excel at capturing a static representation or ‘snapshot’ of a protein, they fall short in depicting dynamic, biologically significant protein movements. Such movements, like the unveiling of cryptic pockets^4^ or allosteric communication^5^, often involve the observation of transient high-energy states, commonly referred to as *rare events*^6,7^

Molecular dynamics (MD) simulations^8,9^ in theory, possess the capability to sample rare molecular events. Yet, a significant limitation emerges; the timescales that MD simulations can access are restricted. As a result, the molecular system frequently remains ensnared in a specific free energy trough. This confinement means it’s challenging to capture these infrequent events with the desired level of detail. Enter a new trick: the stochastic subsampling of multiple sequence alignment (MSA). This technique recently enabled AlphaFold to sample a diverse conformational ensemble of 3D protein structures^10,11^, even those in high energy states. Seeding MD simulations with these AlphaFold generated ensembles followed by Markov State Modelling (MSM) can provide Boltzmann-weighted probability distribution associated with cryptic pocket opening^12^. However, this approach isn’t without its challenges. To gather sufficient samples to construct a reliable MSM^13–15^, one would require a combined simulation time spanning tens of microseconds. Enhanced sampling techniques, especially metadynamics, offer a potent alternative to sample rare transitions within reasonable timescale. Metadynamics accelerates sampling by depositing Gaussian shaped bias along predefined reaction coordinates often known as collective variables^16,17^. Selecting the optimal collective variables^7^, capable of capturing rare transitions, remains at the forefront of biomolecular simulation research.

In this working paper, we introduce slow feature analysis (SFA)^18^, an unsupervised learning algorithm which can capture slowly varying features from high-dimensional temporal data generated by MD simulations. SFA trained on short MD simulations seeded from AlphaFold generated ensemble is used as collective variables in metadynamics simulations to capture cryptic pocket opening and protein-ligand binding in plasmepsin-II^19,20^, a drug target for malaria. We have also shown how SFA can capture *allosteric hotspots* associated with receptor-interacting protein kinase 2 (RIPK2) mediated protein-protein interaction^21^, a key checkpoint associated with inflammatory responses (**Figure 1**). We propose slow feature analysis as a dimensionality reduction algorithm that bridges the gap between AlphaFold and metadynamics and manages to capture Boltzmann distribution associated with a wide range of biological phenomena such as cryptic pocket opening, protein-ligand binding and identifying allosteric hotspots associated with kinase mediated protein-protein interactions.

**Figure 1.**
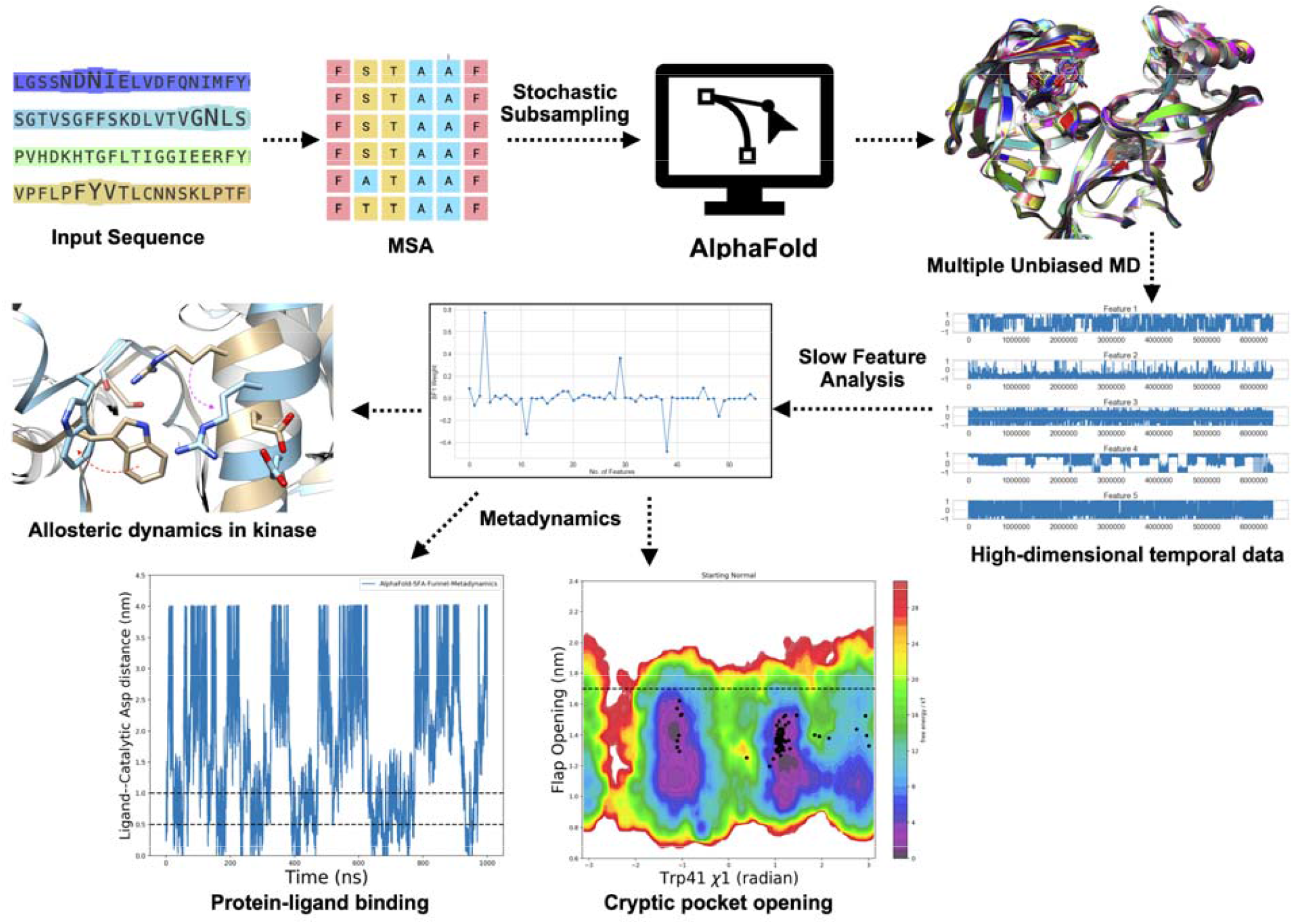
Overview of the pipeline highlighting how to combine AlphaFold, molecular dynamics simulations, slow feature analysis (SFA) and metadynamics to sample *rare events* associated with biomolecular dynamics and molecular recognition. Multiple sequence alignment (MSA) of an input sequence followed by stochastic subsampling enabled AlphaFold to generate a conformational ensemble of plasmepsin II and RIPK2 with structural diversity. AlphaFold generated structural ensemble was used as a starting point to run multiple short unbiased MD simulations which generated high-dimensional temporal data associated with protein dynamics. Slow feature analysis (SFA) captures *slowly varying* features from high-dimensional temporally evolving data. SFA as the collective variables in metadynamics simulations, efficiently sample *cryptic* pocket opening and protein-ligand binding in plasmepsin-II as well as allosteric conformational dynamics in RIPK2.

## Methods

### Structural Ensemble generation using AlphaFold

Structural ensemble for a given sequence was generated using ColabFold implementation of AlphaFold(link:https://colab.research.google.com/github/sokrypton/ColabFold/blob/main/AlphaFold2.ipynb#scrollTo=KK7X9T44pWb7). We generated initial multiple sequence alignment (MSA) using the MMseqs2 method^22^ implemented with ColabFold. We then stochastically subsampled the MSA to a maximum of 32 cluster centers and 64 extra sequences (noted as *max_msa* = 32:64). For the generation of the structural ensemble, we opted for the “*complete*” pairing strategy, which only pairs sequences with a full taxonomic match. We set the number of random seeds to 16 and enabled model dropout. Using dropout, combined with the increased number of random seeds, allows AlphaFold’s neural network to tap into the model’s inherent uncertainties leading to generation of structural ensemble (total 80 structures for plasmepsin-II) with conformational heterogeneity.

Conformational ensemble of RIPK2 was generated using an older version of ColabFold (https://colab.research.google.com/github/sokrypton/ColabFold/blob/main/beta/AlphaFold2_advanced.ipynb). A structural ensemble consisting of 32 structures of RIPK2 were generated using the following settings as described previously: *msa_method: MMseqs2, pair_mode=unpaired+paired, pair_cov=25, pair_qid=20, max_msa=32:64, subsample_msa=True, num_models=1, num_samples=32, num_ensemble=1, max_recycles=3, is_training=True*.

### Molecular Dynamics Simulations

Each structure from the conformational ensemble generated by AlphaFold was prepared for molecular dynamics simulations using the *tleap* module from Amber2022^23^, following the protocol outlined by *Meller et al*^12^. In brief, the proteins were parameterized with the AMBER FF14SB force field^24^. To achieve system neutrality, 17 Na^+^ ions were added to each system. Systems were then solvated within a truncated octahedron box, ensuring a minimum of 10 Å between the protein and the edge of the box. The system underwent a two-phase minimization: (a) an initial phase where only the water and ions were minimized while the protein was held in place using a restraint potential of 100 kcal/mol^−1^ Å^2^ (200 steepest descent steps followed by 200 conjugate gradient steps), and (b) an unrestrained minimization of the entire system over 500 steps.

After the minimization process in Amber2022, we transformed Amber topologies into Gromacs format via Acpype^25^. Each system was gradually heated from 0 to 300 K for 500 ps in an NVT ensemble, with harmonic restraints (500 kJ mol^−1^nm^−2^) on the backbone’s heavy atoms. Subsequently, systems were equilibrated for 200 *ps* in an NPT ensemble at 300 K, devoid of restraints. The *Parrinello–Rahman* barostat^26^ ensured a consistent pressure of 1 bar, while the v-rescale thermostat controlled the temperature. Production runs were performed in the NPT ensemble, maintaining conditions at 300 K and 1 bar. The leapfrog integrator and Parrinello– Rahman thermostat was employed with a 2 fs timestep. Nonbonded interactions had a cutoff of 1.0 nm and long-range electrostatic interactions were treated using the Particle Mesh Ewald (PME) method^27^ with a 0.16 nm grid spacing. The LINCS algorithm^28^ was used to constrain H-bonds. Heating, equilibration, and production runs were performed using Gromacs 2022^29^.

For the 80 structures of plasmepsin-II produced by AlphaFold, we conducted two independent 40ns production runs, each with unique initial starting velocities. For 32 structures of RIPK2, we performed ten independent 20 ns productions runs each with different initial velocities. Trajectories were saved every ps. Molecular dynamics simulation of RIPK2+XIAP complex (PDB: 8AZA) was also performed using a similar protocol. The missing residues were modelled using Modeller web server^30^ integrated with UCSF Chimera.

### Slow Feature Analysis (SFA)

Slow Feature Analysis (SFA) is a dimensionality reduction technique designed to process high-dimensional, temporally evolving data. Its primary goal is to transform a J-dimensional input signal, (*c*(*t*)), using a set of nonlinear functions,*g*_*k*_(*c*), to produce output signals *y*_*k*_(*t*)= *g*_*k*_(*c*(*t*)) These output signals are optimized to minimize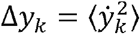, where 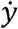 represents the derivative of *y*, and ⟨ ⟩ indicates temporal averaging. This minimization targets the extraction of features that vary slowly over time. SFA also imposes additional constraints as follows:

a. each output feature must have zero mean ⟨ y_k_ ⟩ =0,
b. unit variance 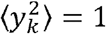, and
c. decorrelated from others ⟨ *y*_*k*_ *y*_*k*_′ ⟩ =0 for all *k*′ < *k*

These constraints ensure that each extracted feature is scaled similarly, uncorrelated with others, and avoids the trivial solution, *y*_*k*_ =*c* where *c* is a constant. When detailing the algorithm below, signals represented by capitals represent raw signals, while signals represented by lower-case represent normalized signals.

To perform the SFA algorithm, first start with a J-dimensional input signal *X*(*t*). Next, normalize the input signal to get:

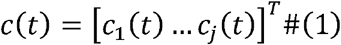

where:

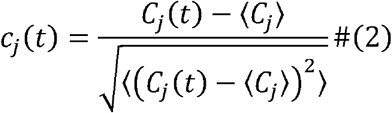

Such that:

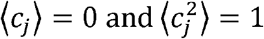

Next, perform a linear or a non-linear expansion using a set of functions *H*(*c*) to produce an expanded signal *Z*(*t*). In the case of a quadratic expansion this would include monomials of degree one and degree two including mixed terms as shown below, which would result in quadratic SFA:

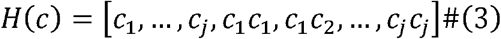

and thus:

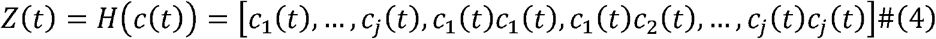

The next phase involves sphering (or whitening) of *Z*(*t*), generating a normalized signal *Z*(*t*) through the equation:

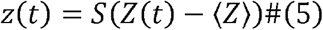

where:

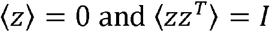

In this case,*I* is the identity covariance matrix and *S* is a sphering matrix. Matrix *S* can be solved by performing PCA on matrix (*Z*(*t*) − ⟨*Z*⟩)

Further, PCA is applied to the matrix ⟨*Ż Ż*^*T*^⟩ where *Ż* is the time derivative of the sphered expanded signal *Z*(*t*). The *K* eigenvectors with the lowest eigenvalues λ_*k*_ result in the normalized weight vectors *w*_*k*_ satisfying:

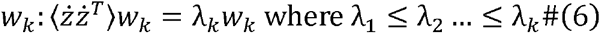

This leads to the formulation of the desired set of real-valued functions *g*(*c*) =[*g*_1_ (*c*),…, *g*_*k*_ (*c*)]^*T*^ where:

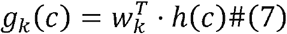

and:

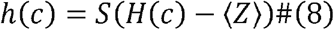

Here, *S* is the sphering matrix that was earlier solved to normalize the expanded signal *Z*(*t*) Now, define the output signal *y*(*t*)as:

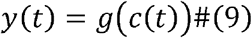

where:

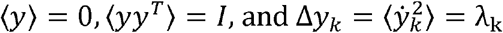

In this formulation, the components of y(t), representing the extracted slow features, are characterized by zero mean, unit variance, and mutual decorrelation, thereby encapsulating the core principles of SFA.

### Application of SFA on molecular dynamics

Slow feature analysis was performed on training data generated from unbiased molecular dynamics simulations launched from the conformational ensemble of plasmepsin II (80 structures * 2 clones each * 40 ns = 6.4 microsecond) and RIPK2 (32 structures * 10 clones each * 20 ns = 6.4 microsecond) generated by AlphaFold.

In this study, we adopted linear Slow Feature Analysis (SFA) for our experimental analysis, implemented in the *sklearn SFA* package (https://sklearn-sfa.readthedocs.io/en/latest/). Our initial step involved the compilation of featurised trajectory data into a J-dimensional input signal, denoted as *c*(*t*) and normalizing the data by solving for the whitening matrix *S* for the input signal using PCA and subsequently transforming it into c^*white*^. Whitening can be expressed as the linear map 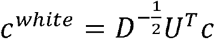 where the covariance matrix of *c*(*t*) is decomposed as *C*_*c*_=*UDU*^*T*^

We then do PCA on finite differences of the whitened input signal 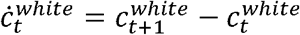 to get the decomposition of the covariance matrix for the finite differences *ċ* such that 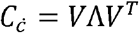.

The eigenvectors corresponding to the smallest eigenvalues in this decomposition represented the normalized weights of the slowest moving features. These extracted weights, derived from the training data, were then utilized to construct a new collective variable for subsequent metadynamics simulations. This approach leverages the strength of linear SFA in distilling critical dynamical features from high-dimensional data, enabling nuanced exploration and modeling of the underlying processes.

### Markov State Model

Markov state model was performed on sin and cos transformed □1 and □2 angles of Trp41 extracted from unbiased molecular dynamics simulations (80 structures * 2 independent clones * 100ns each = total 16 μs) launched from AlphaFold generated conformational ensemble of plasmepsin-II. *Kmeans* clustering was performed on the transformed dihedral space with *k=200*. Finally, maximum likelihood MSM was generated using a lag time of 6 ns (Figure S2 highlights the implied timescale plot). Equilibrium populations extracted from MSM is projected along different features to highlight conformational heterogeneity associated with plasmepsin II. PCCA+^31^ was used to generate macrostate definition which manages to distinguish conformational states associated with Trp41 (Figure S2). MSM generation was performed using PyEMMA 2.5.7^32^.

### Metadynamics

Well-tempered metadynamics simulations were performed using the first two slow features as collective variables (CVs) at 300K. For plasmepsin-II, gaussians of height 1.50 kJ/mol were deposited at every 500 steps. The Gaussian widths for the first two slow features were set at 0.32 and 0.25, determined by taking approximately one-third of the standard deviation from the unbiased training data. The bias-factor was set at 20. We performed two distinct metadynamics simulations, each lasting around ∼400 ns. One began from the closed state (PDB: 1LF4)^33^ and the other from the cryptic pocket open state (PDB: 2BJU)^34^. For RIPK2, 500 ns well-tempered metadynamics simulation was performed using first two slow features as CVs using a gaussian widths of 0.16 and 0.25, height 1.50 kJ/mol and bias factor 20. Metadynamics simulations were performed using Gromacs 2022 patched with Plumed 2.7^35^. Unbiased free energy surfaces along different features were extracted using the reweighting protocol developed by *Tiwary and Parrinello*^36^.

Funnel metadynamics^37,38^ represents an enhanced sampling technique designed to improve sampling efficiency of protein-ligand binding. This method specifically targets the sampling process along the pathway of ligand unbinding/binding. It employs a unique, funnel-shaped restraint potential, effectively narrowing down the exploration space when the ligand is in its unbound state leading to rapid binding/unbinding of ligand. Funnel metadynamics simulations with and without SFA was performed using a small molecule bound to ‘cryptic pocket’ of plasmepsin II (PDB: 7QYH). The funnel potential is highlighted in **Figure 2**. The following parameters were used to define the funnel: *Z*_*cc*_=2.0 nm, *R*_*cyl*_ = 0.1 nm, and α = 0.5 radian.

**Figure 2.**
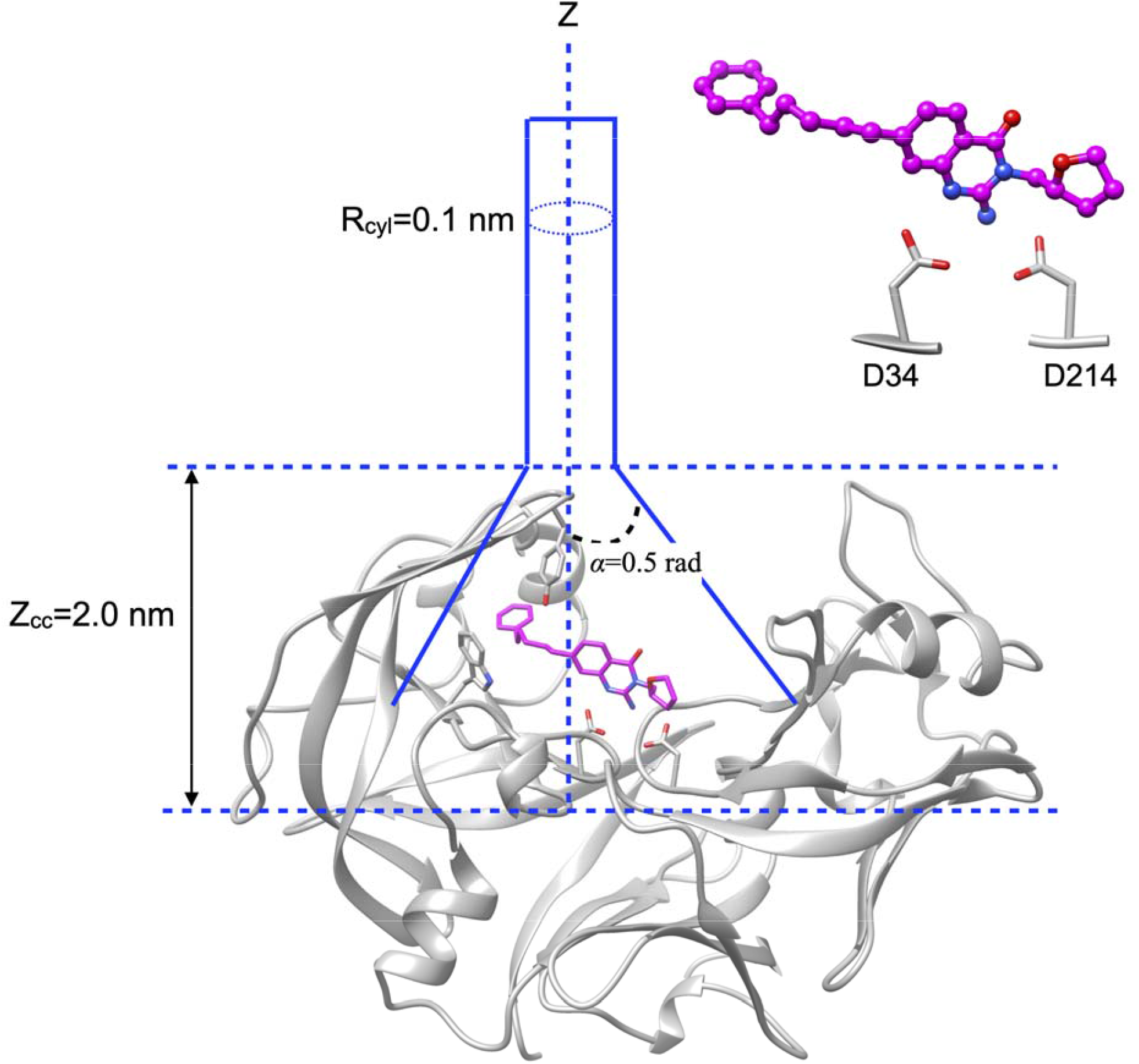
A visual depiction of the funnel setup employed in metadynamics simulations, emphasizing the definition of various key parameters integral to these simulations. Central to this setup is the measurement of the distance between the center of mass of the catalytic aspartate dyads (specifically D34 and D214) and the ligand. This serves as the ligand binding/unbinding collective variable (CV) in traditional funnel metadynamics. A small molecule (highlighted in *magenta*) bound to the cryptic pocket of plasmepsin II (PDB: 7QYH) was used for funnel metadynamics simulations.

Traditional funnel metadynamics was performed using the distance between the center of mass (COM) of ligand and Cγ atoms of catalytic Asp dyads (Asp 34 and 214) as CV. A steep repulsive wall was applied along the COM distance with a spring constant of 50000 kJ/mol/nm^2^. Other details regarding the funnel setup have been described in the Supporting information. Finally, well-tempered metadynamics was performed using a Gaussian height of 1.5 kJ/mol, width of 0.03 nm and bias factor of 20. SFA-Funnel metadynamics was performed using first two slow features as additional CVs as described in the previously. Funnel metadynamics in principle can predict protein-ligand binding free energy. In our study, given the lack of experimental binding free energy data, we aimed to determine whether SFA-augmented funnel metadynamics can accelerate ligand binding.

## Results

### SFA captures critical fluctuations necessary for cryptic pocket opening

SFA trained on small independent unbiased molecular dynamics simulation launched from a structural ensemble generated by AlphaFold managed to capture flipping of Trp41 in plasmepsin II, necessary for cryptic pocket opening. It also captured flipping of Tyr77 along □1 angle as another key feature (Figure S3). Flipping of Tyr77 in conjunction with flipping of Trp41 exposes a fully ‘open’ cryptic pocket primed for ligand binding.

A recent study highlighted how microsecond long unbiased MD simulation launched from apo plasmepsin II failed to capture cryptic pocket opening^12^. To test the effectiveness of SFA as CVs with metadynamics we performed two independent simulations, one launched from ‘closed’ state (PDB: 1LF4) and the other from the cryptic pocket ‘open’ state (PDB: 2BJU, removing the ligand to make it apo). It is key to highlight unbiased MD simulations launched from *apo-like* ‘open’ state failed to sample open ↔ closed transitions (Figure S1, Supporting information).

Metadynamics simulations using first two slow features as CVs managed to sample multiple flipping events along Trp41 □1 and □2 angles leading to close ↔ open transitions in plasmepsin II within ∼350-400 ns of simulation time. Reweighted free energy surfaces from metadynamics simulations agreed extremely well with MSM suggesting that we reached convergence (**Figure 3**) for the choice of force field and water model (see Figure S4 and S5 in Supporting Information regarding convergence of metadynamics simulations).

**Figure 3.**
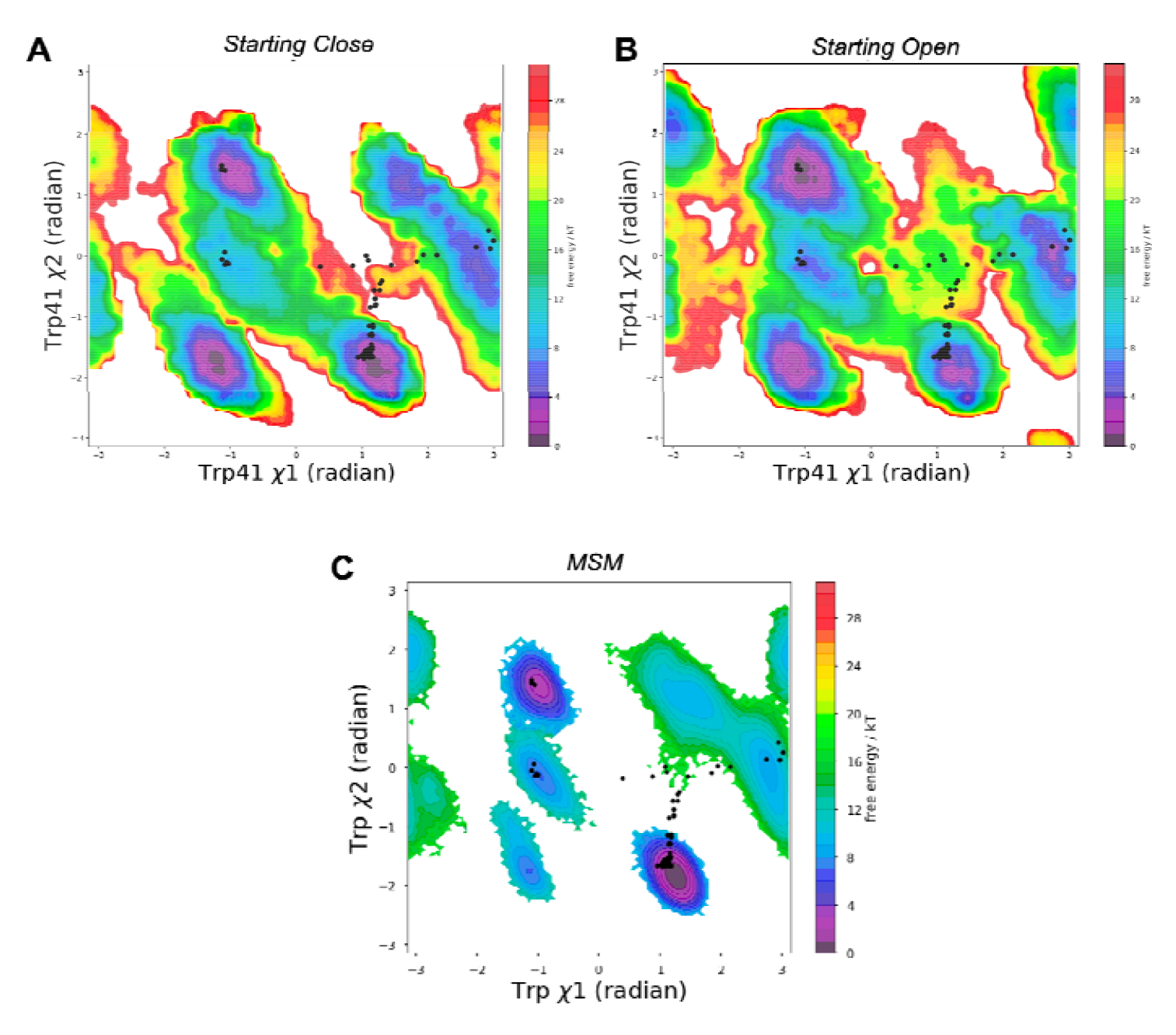
Metadynamics simulations using first two slow features as CVs managed to capture Trp41 flipping necessary for cryptic pocket opening in plasmepsin II. Reweighted free energy surface from metadynamics, starting from close (A) and open (B) states of plasmepsin II yields a similar landscape when compared with Markov state model approach (C). It is important to note that the simulation length for the metadynamics simulations is ∼400ns each whereas the Markov state model was performed using an aggregate of 16 μs of simulation data. AlphaFold generated structural ensembles are highlighted in *black dots*.

Metadynamics also captured flipping of Tyr77 (Figure S6 in Supporting Information) which is a key residue governing the flap ‘opening’ in plasmepsin II. Opening of the flap in conjunction with Trp41 flipping exposes the ‘deep’ cryptic pocket (**Figure 4**).

**Figure 4.**
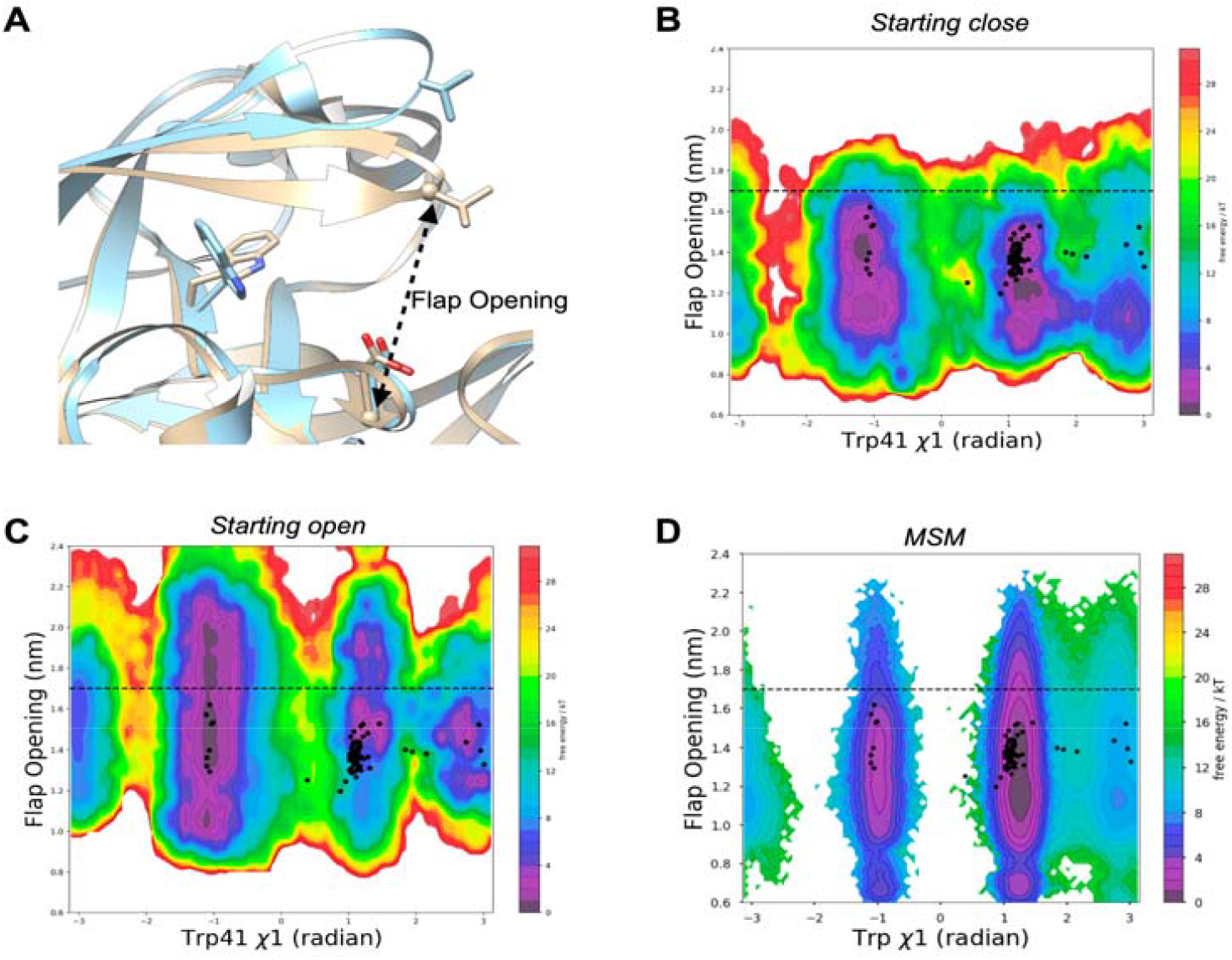
SFA-metadynamics managed to capture a deep cryptic pocket opening in plasmepsin II. (A) Flap opening is defined by the *C*α*--C*α distance between Asp34 and Val78. Flap opening in conjunction with Trp41 flipping (blue) exposes a deep cryptic pocket in plasmepsin II when compared with close apo conformation (silver). (B, C) Metadynamics simulations starting from close and open apo-like states managed to sample ‘deep’ cryptic pocket opening (Flap opening distance > 1.7 nm) within ∼400ns when compared with MSM (total ∼16μs of aggregate simulation). It is important to highlight that the AlphaFold generated ensemble (black dots) failed to sample a deep cryptic pocket opening in plasmepsin II.

Metadynamics using SFA CVs also managed to capture an alternate flipped state of Trp41 which has not been sampled by AlphaFold. Such a state has been captured by holo crystal structure of plasmepsin II, PDB: 4Z22^39^ (Figure S7 in Supporting information).

### SFA augmented funnel metadynamics accelerates ligand binding in plasmepsin II

Ligand binding in the cryptic pocket of the plasmepsin-II requires correct orientation of Trp41 as well as opening of the flap which allows ligand entry. We performed two sets of funnel metadynamics: a) traditional funnel metadynamics with distance between center of mass of the ligand and catalytic aspartic acid dyads as a CV and b) SFA augmented funnel metadynamics which uses first two slow features trained on AlphaFold seeded MD simulations as orthogonal CVs in conjunction with distance CV (see Methods).

SFA augmented funnel metadynamics managed to capture multiple ligand binding events compared to traditional funnel metadynamics. Moreover, SFA-funnel metadynamics captures opening of the flap as well as flipping of Trp41, two key conformational events require for ligand binding (**Figure 5**). Previously *Bobrovs and co-workers*^40^ highlighted the necessity of using dynamical information associated with flap opening via ‘path’ as orthogonal CVs to capture accelerated ligand binding when compared with traditional funnel metadynamics. SFA trained on AlphaFold seeded MD simulations captures flipping of Tyr77, key residue governing flap opening in plasmepsin II. SFA-funnel metadynamics not only corroborates the findings of *Bobrovs and co-workers* but also significantly underscores the integral role of protein dynamics and structural adaptability in the process of ligand binding.

**Figure 5.**
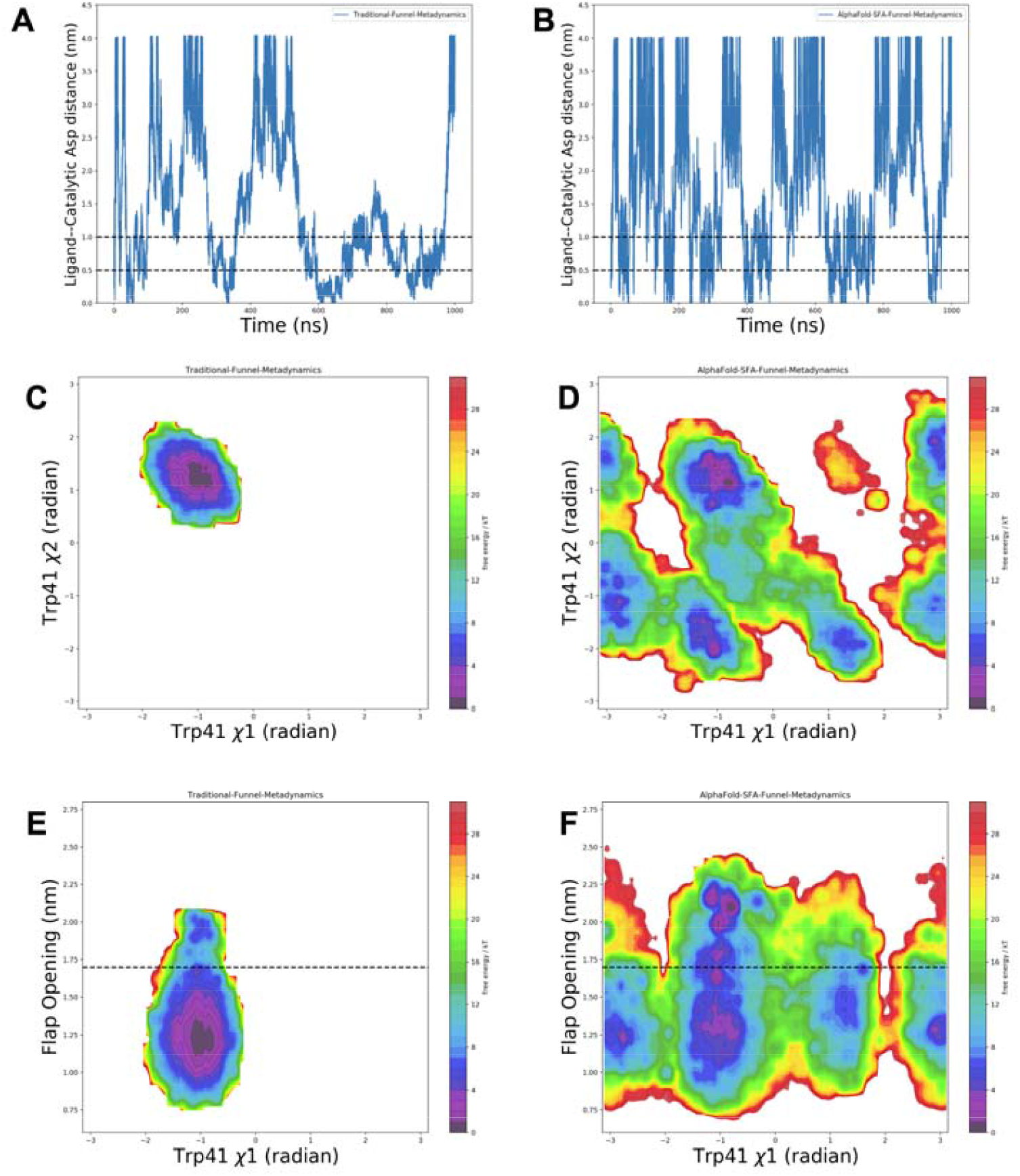
SFA augmented metadynamics using distance between ligand and the COM of the catalytic aspartic acids as binding/unbinding CV, resulted in more frequent observations of ligand binding compared to traditional funnel metadynamics (A, B). Ligand binding in the deep cryptic pocket is indicated by a ligand-catalytic Asp distance of less than 0.5 nm. A distance greater than 0.5 nm but less than 1.0 nm suggests the ligand is in an intermediate region, maintaining contact with the protein. A distance exceeding 1.0 nm implies the ligand is completely unbound from the protein. Additionally, incorporating slow features as orthogonal CVs in funnel metadynamics successfully captured two rare conformational transitions: the flipping of Trp41 (C, D) and the opening of the flap (E, F), both of which are vital for ligand binding.

**Figure 6.**
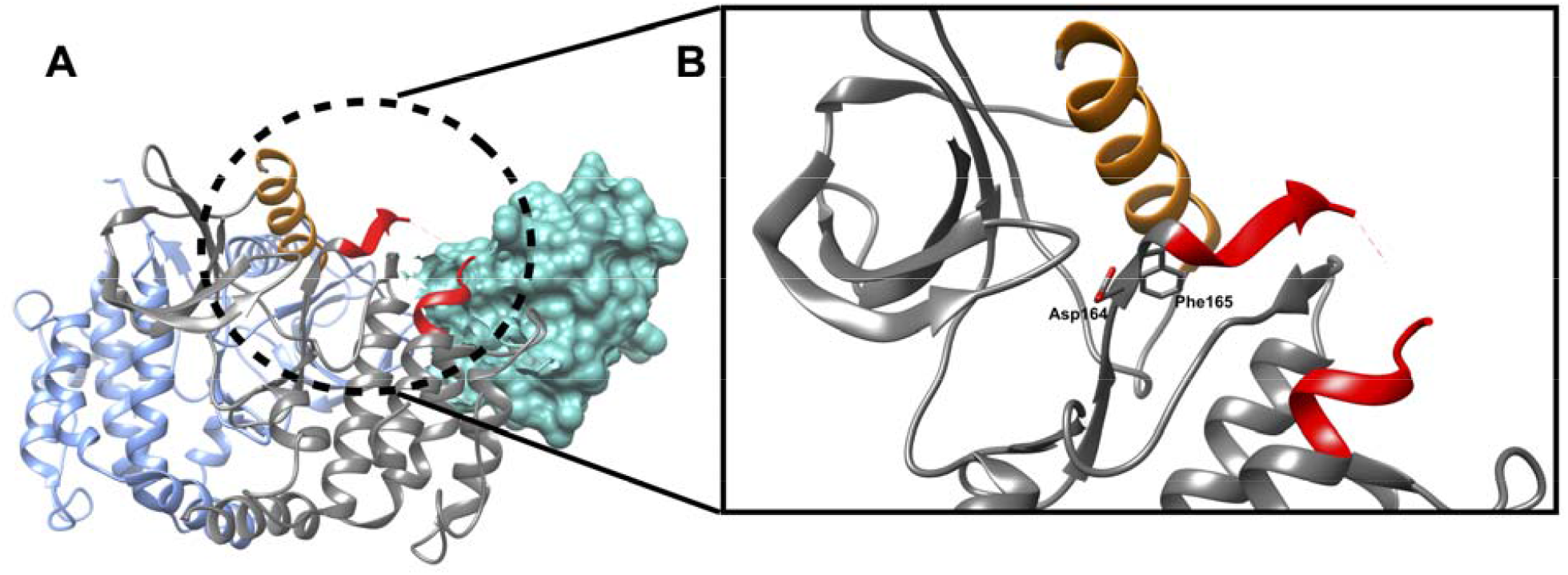
**A:** Cryo-EM structure (PDB: 8AZA) of RIPK2 and XIAP. In this structure, XIAP is visualized with a sea green surface and is located away from the RIPK2’s active site. **B:** The active site of RIPK2, characterized by the DFG motif (Asp164, Phe165, Gly166), engages with XIAP through the kinase’s activation loop, depicted in red. Notably, sections of the activation loop remain unresolved in both cryo-EM and X-ray structures of RIPK2, indicating dynamic nature of the region. The αC helix, adjacent to the DFG motif and highlighted in orange, is posited to significantly influence RIPK2’s conformational changes by its interactions with the activation loop. However, the precise mechanisms by which the αC helix and activation loop affect RIPK2’s conformation and its interaction with XIAP are not yet fully understood and remain an *open question*.

### SFA captures allosteric hotspots in RIPK2

Receptor-Interacting Protein Kinase 2 (RIPK2) plays a crucial role in immune and inflammatory responses. It functions as a key signaling molecule in the NOD-like receptor pathways, which are vital for recognizing intracellular pathogens and danger signals. Upon activation, RIPK2 undergoes ubiquitination, a process significantly facilitated by the E3 ubiquitin ligase XIAP (X-linked Inhibitor of Apoptosis Protein). This interaction between RIPK2 and XIAP is pivotal in modulating downstream signaling pathways, especially the NF-κB and MAPK pathways, which are essential for the production of pro-inflammatory cytokines and immune responses. The RIPK2-XIAP interaction thus represents a critical checkpoint in the regulation of inflammation and innate immunity, highlighting its potential as a therapeutic target in diseases characterized by excessive or chronic inflammation^21,41^. The interaction between RIPK2 and XIAP is intricately regulated by the activation loop and allosteric modulation in RIPK2^42^. The activation loop, a part of RIPK2’s kinase domain, undergoes conformational changes upon activation, influencing its interaction with XIAP via allostery. However, due to its dynamical nature, the activation loop in RIPK2 often remain unresolved in X-ray crystallography, posing challenges in comprehending how the activation loop’s conformational diversity allosterically influences its interaction with XIAP.

SFA trained on AlphaFold seeded MD simulation managed to capture several critical conformational fluctuational associated with RIPK2 (**Figure S8**). One of the key structural features captured by SFA is the flipping of Phe165 in the DFG moiety. Metadynamics simulation using first two slow features as CVs managed to capture Phe165 flipping in apo RIPK2, a rare event which has not been sampled in ∼500 ns unbiased MD simulation (**Figure 7, 8**). It is important to highlight that such flipping has only been observed (**Figure S11**) in holo RIPK2 bound with small molecules, GSK583 (PDB: 5J7B^43^), SB-203580 (PDB: 5AR4^44^), pyrazolocarboxamide scaffold (PDB: 6SZJ^45^) and CSLP18 (PDB: 6FU5). SFA also managed to capture flipping of Trp170, a key residue in present in the activation loop acting a hallmark of allostery mediated conformational transition in RIPK2. SFA-metadynamics allowed us to sample a high-resolution picture of the allosteric hotspot involving Trp170. In the active state of RIPK2, Trp170 points towards αC helix which is represented by distance between Glu68-CD—Trp170-NE1. Another structural feature of the active RIPK2 is the Arg65 sidechain mediated H-bond interaction involving Ser168 (captured by the distance between Arg65 and Ser168) (**Figure S12**). This H-bond interaction keeps the stabilized the activation loop which facilitates interaction with XIAP (**Figure S14**). SFA-metadynamics managed to capture rare transition between active <—> inactive states of RIPK2. A hallmark of such transition is the flipping of Trp170 which resulted in the breaking of H-bond interaction involving Arg65 (**Figure 7, 8)** which shifts the αC helix in an ‘outward’ conformation.

**Figure 7.**
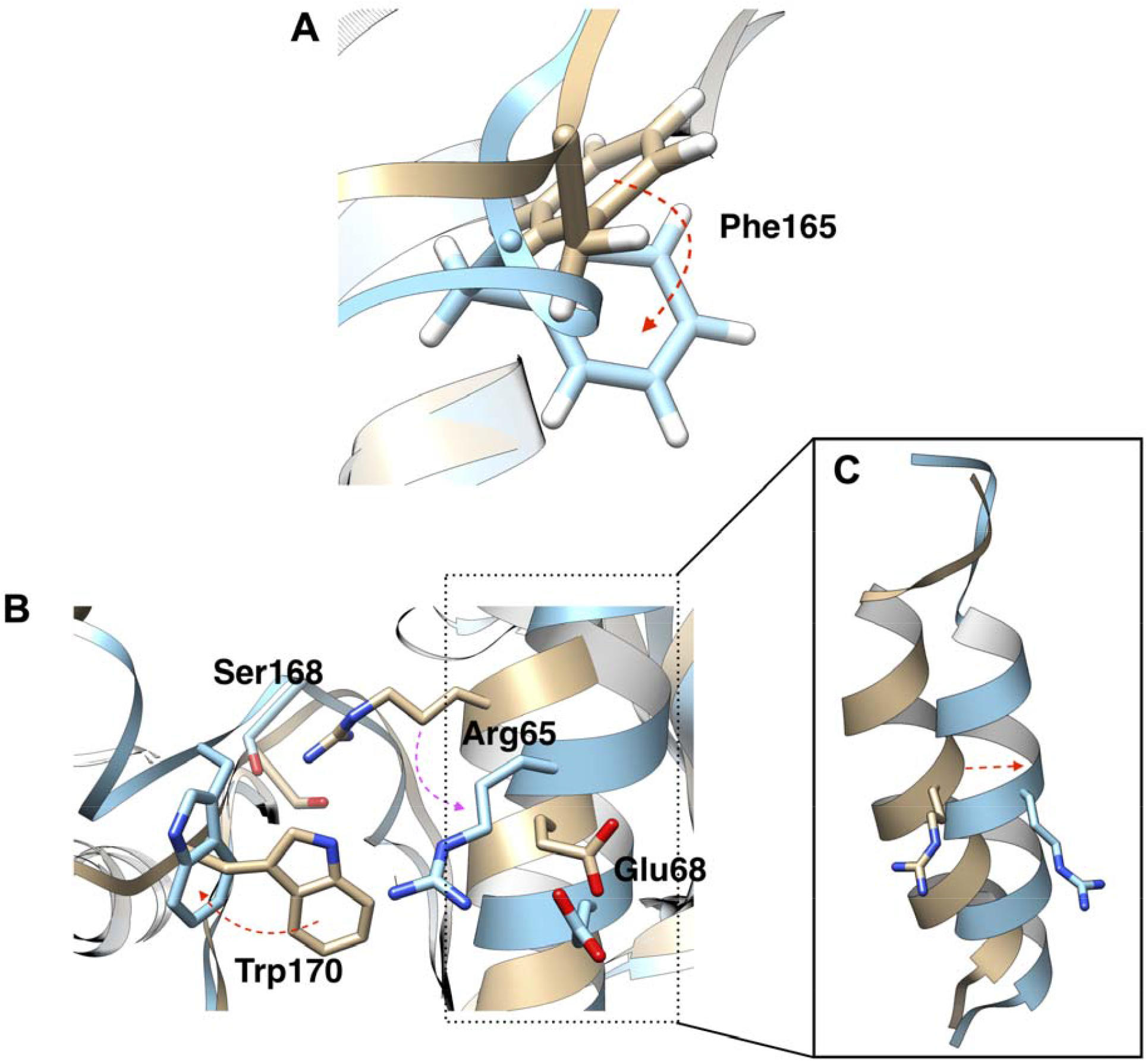
SFA metadynamics has successfully delineated the intricate conformational dynamics underlying the active to inactive transitions in RIPK2. A pivotal structural element in this transition is the flipping of Phe165 within the DFG motif (A). Our SFA-metadynamics approach not only captured multiple instances of the Phe165 flip but also effectively sampled additional critical transitions. These include the flipping of Trp170 and the disruption of the hydrogen bond between Arg65 and Ser168 (B). Such molecular events lead to the displacement of the αC helix into an ‘outward’ orientation (C), a hallmark of the inactive state of RIPK2.

**Figure 8.**
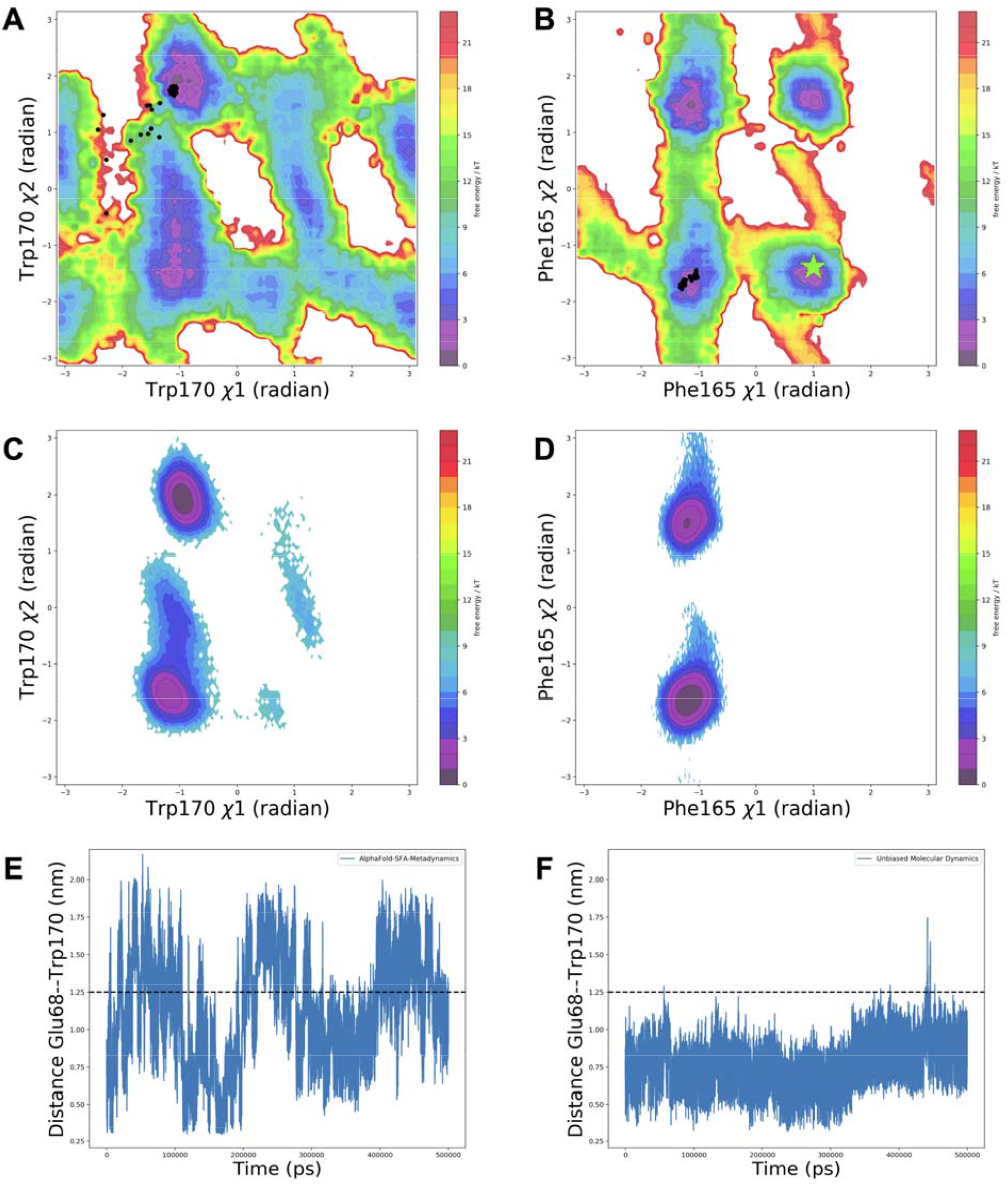
Comparative analysis of the conformational sampling in RIPK2 using SFA-metadynamics and traditional unbiased MD simulation. The upper panel (A and B) showcases the reweighted free energy surface associated with 1 and 2 angles of Trp170 and Phe165 (see Figure S10 for convergence of SFA-metadynamics along reweighted FES), highlighting accelerated sampling achieved through SFA-metadynamics. The black dots highlight values of dihedral angles in AlphaFold generated conformational ensemble. The ‘*green star’* indicates the value of dihedral angles in holo RIPK2 (GSK583-RIPK2 complex, PDB: 5J7B). The middle panel (C and D) shows population distribution of the aforementioned CVs in 500ns unbiased MD simulation which failed to sample such conformational transitions. The bottom panel (E and F) illustrates the time-trace of distance between Trp170 and Glu68, a key indicator of active to inactive transition in RIPK2. A demarcated line at 1.25 nm distinguishes the active (distance < 1.25 nm) and inactive states (distance > 1.25 nm) of RIPK2, underscoring the critical distance for the transition. These CVs highlights a stark contrast between the two methodologies, emphasizing the superior ability of SFA-metadynamics to capture multiple key conformational transitions that are pivotal for the RIPK2’s functional dynamics.

Such conformational transition destabilizes the activation loop which blocks XIAP binding. Molecular simulation of XIAP-RIPK2 complex highlighted a critical H-bond interaction involving Arg171 of RIPK2 and Asp214 of the XIAP (**Figure S14, S15**). This interaction is dependent on the stabilization of the activation loop involving Trp170 and Arg65. Overall, our study highlighted how SFA trained on AlphaFold seeded molecular simulation augmented by metadynamics captures previously illusive allosteric hotspot involving activation loop of RIPK2 which governs RIPK2+XIAP interactions.

## Discussion

Stochastic subsampling of MSA allowed AlphaFold to sample a structural ensemble of plasmepsin II with conformational diversity; however, it failed to sample the ‘deep’ cryptic pocket opening (Figure 3). Well-tempered metadynamics simulations with first two slow features as CVs managed to capture ‘deep’ cryptic pocket opening in plasmepsin II within a factor of total simulation length when compared to MSM based approach. Deep cryptic pocket opening is a rare event in plasmepsin II which is a combination of flipping of Trp41 and flap opening. SFA-augmented funnel metadynamics outperformed traditional funnel metadynamics in sampling the complex dynamics of ligand binding and unbinding to the cryptic pocket of plasmepsin II. This approach successfully captured multiple recrossing and enabled more effective conformational sampling. Specifically, it facilitated the sampling of crucial conformational transitions like the flap opening and the flipping of Trp41, both vital for ligand binding. In contrast, traditional funnel metadynamics without SFA did not adequately sample these transitions, notably failing to capture the flap opening and Trp41 flipping, thus impeding effective ligand binding. Additionally, in the absence of SFA, traditional funnel metadynamics showed that ligand unbinding led to formation of a hydrogen bond formation between Tyr77 and Asp34, as noted by Bhakat and Söderhjelm^20^, which obstructed flap opening. Remarkably, SFA, when trained on AlphaFold-seeded MD simulations, was able to detect the flipping of Tyr77, a critical movement for enabling flap opening. Utilizing the first two slow features as collective variables in funnel metadynamics significantly enhanced the sampling efficiency of flap opening, thereby promoting ligand binding.

Using AlphaFold-seeded MD simulations, SFA effectively identifies allosteric hotspots in RIPK2. These hotspots are crucial for the interaction between RIPK2 and XIAP, a key element in NOD1/2-mediated immune responses, with implications in inflammatory bowel disease and other conditions. The study reveals how the conformational dynamics of the activation loop and the Asp-Phe-Gly (DFG) loop in RIPK2 regulate active <—> inactive transition (**Figure 9**). This transition involves intricate dynamics associated with the αC helix, DFG loop, activation loop, and other distant residues.

**Figure 9.**
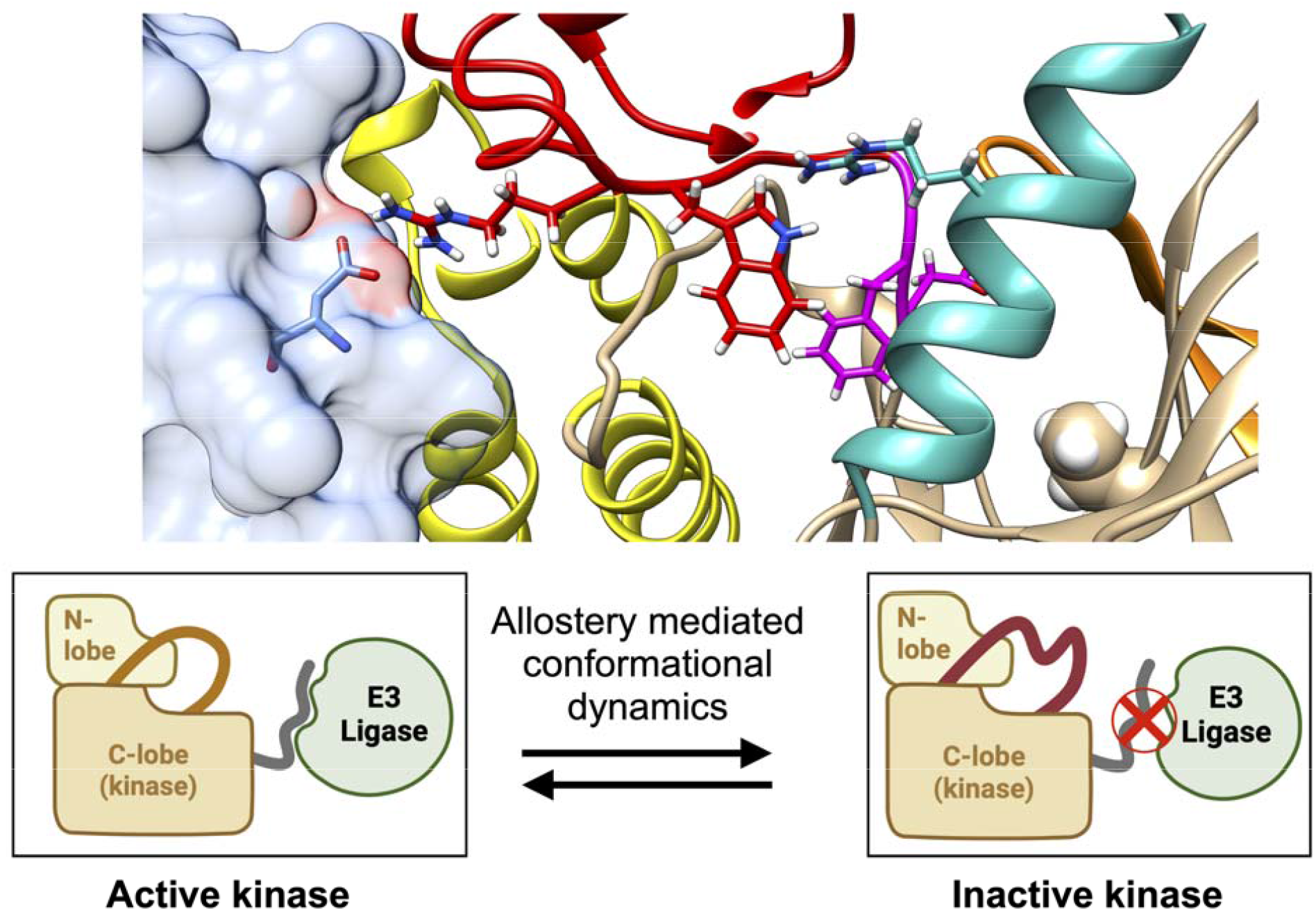
Illustration of the novel mechanism linked to the conformational dynamics of RIPK2, emphasizing the shift between its active and inactive states. In its active form, RIPK2 engages in PPI interactions with the E3 ligase, XIAP, while the inactive state prevents these interactions. Critical residues within the DFG loop (magenta), activation loop (red), and αC helix (sea green) regulate the transition between active and inactive states, thereby influencing the interactions between RIPK2 and XIAP (blue surface).

SFA-augmented metadynamics successfully captures two vital interactions: 1) hydrogen bond interactions between the Arg65 sidechain and Ser168, monitored by the distance between Arg65 and Ser168, and 2) the interaction involving Trp170 and residues in the αC helix, where Trp170’s side-chain NH group is oriented towards Glu68, measured by the distance between Trp170 and Glu68. These observations build upon Pellegrini et al.’s report of a water-mediated hydrogen bond between Trp170 and Glu68^46^. These interactions are pivotal in defining RIPK2’s *active state*, stabilizing the activation loop, which serves as a binding platform (*‘landing pad’*) for XIAP. Notably, the H-bond interaction of Arg171 in the activation loop with Asp214 in XIAP is essential for stabilizing the RIPK2-XIAP complex.

The transition to the *inactive state* involves the disruption of these interactions, increased conformational entropy of the activation loop, and the flipping of Trp170. This flipping breaks the interactions between Trp170 and the αC helix, as well as between Arg65 and Ser168, thus destabilizing the activation loop. Concurrently, the flipping of Phe165 in the DFG loop shifts the αC helix outward and increases the mobility of the activation loop, dismantling the structural integrity of the ‘*landing pad*,’ impeding XIAP binding. Careful analysis of another serine/threonine kinase, BRAF also highlights the coupled motion involving DFG-Phe *flipping* and ‘*outward*’ conformation of αC helix (**Figure S16**). Additionally, SFA highlights the significance of the flipping motions of Ile208 and Lys209 in RIPK2’s regulatory region. Lys209 forms crucial interactions with Glu211 (**Figure S17**) in XIAP. The flipping of Ile208 and Lys209 disrupts these interactions, contributing to the inhibition of XIAP binding. This aligns with experimental findings that indicate the mutation of Lys209 impedes the RIPK2-XIAP interaction^42^.

A careful examination of 3D structures of small molecules bound to RIPK2, as deposited in the Protein Data Bank (PDB), reveals two distinct categories: (a) structures in which the position of Trp170 remains unresolved, and (b) structures where Trp170 is oriented towards Glu68 (hallmark of active RIPK2). Notably, in structures where Trp170’s position is indeterminate, Phe165 appears in a ‘*flipped*’ state, as evidenced in PDB entries 5J7B, 5AR4, 6SZJ and 6FU5. We propose that small molecules binding to these specific PDB structures function as RIPK2-XIAP inhibitors, potentially inducing an allosteric conformational change as previously discussed. This insight paves the way for the rational design of novel RIPK2-XIAP inhibitors, targeting inflammatory diseases and beyond.

Recent works highlighted how we can combine structural ensembles generated by AlphaFold with autoencoder framework^47^ and MSM^12^ to predict Boltzmann distribution and capture rare events. The enhanced sampling strategy we propose, termed SFA-metadynamics, augments the toolkit of approaches aiming to extract the Boltzmann distribution from structural ensembles generated by AlphaFold. This strategy is particularly effective to capture rare transitions, such as opening of the ‘deep’ cryptic pocket in plasmepsin II. SFA in principle is similar to time-structure based independent component analysis (tICA)^48^. However, SFA is based on a straightforward idea: extract uncorrelated output signals that are ordered by slowness. This allows SFA to capture slowly varying features from high-dimensional and noisy temporal data generated by AlphaFold seed molecular dynamics simulations. SFA can also act as a reduced space on which one can build MSM. Further, conformations which contribute to the first few slow features can be used as seeds to launch small MD simulations. These MD simulations can be stitched together using MSM to predict thermodynamics and kinetics associated with conformational transitions. Kinetics extracted from MSM built on top of SFA can be directly compared with infrequent metadynamics^49^ with slow features as CVs. In essence, SFA emerges as an innovative technique that bridges the gap in the estimation of kinetics linked to rare events, whether one uses MSM or metadynamics.

## Conclusions

In this study, we present a novel integration of slow feature analysis (SFA), derived from molecular dynamics (MD) simulations initiated by AlphaFold predictions, with metadynamics. This innovative approach is applied to sample cryptic pocket opening, protein-ligand binding/unbinding, and the allosteric dynamics within biomolecules. By synergistically combining SFA with metadynamics, we successfully identified multiple transitions between the closed and open states of the cryptic pocket in plasmepsin II in just a few hundred nanoseconds. This is a significant improvement over traditional methods, which typically require several microseconds of simulation data to develop a comprehensive Markov state model. Furthermore, the SFA augmented funnel metadynamics not only effectively captured multiple instances of ligand binding and unbinding but also explored essential conformational dynamics of protein that facilitate ligand binding. Additionally, our SFA-metadynamics approach discovered the pivotal role played by the activation loop and the DFG moiety in the transition from the active to inactive state in RIPK2. This transition is a critical molecular event in the interaction between RIPK2 and XIAP, which has implications in the pathogenesis of inflammatory diseases.

This work underscores a novel approach that merges the capabilities of AlphaFold, SFA, and metadynamics. It offers a robust framework to predict the Boltzmann distribution corresponding to conformational shifts necessary to capture rare events such as cryptic pocket opening, protein-ligand binding, and allosteric modulation in biomolecules. Our approach identified crucial conformational alterations linked to apo RIPK2 that allosterically regulate the interactions between RIPK2 and XIAP. This forms the basis for our forthcoming research, which will explore how ligand binding affects RIPK2-XIAP interactions by inducing a shift in the population between RIPK2’s active and inactive states. It is important to highlight that in order for methods such as SFA, deep-TICA^50^, RAVE^51^ to be useful, the training data should transiently sample conformational heterogeneity. Short molecular dynamics simulations launched from AlphaFold generated ensemble encompasses necessary conformational heterogeneity which can be extracted by SFA and used as CVs within enhanced sampling framework such as metadynamics to efficiently sample rare conformational transitions which plays a key role in molecular recognition and conformational dynamics of biomolecules. Here, we develop AlphaFold-SFA which bridges the gap between AI based protein structure prediction model, molecular dynamics simulation and metadynamics to sample three different rare events of biological relevance, highlighting the power of our method in structure-based drug discovery.

## Supporting Information

Supporting information contains population distribution in unbiased molecular dynamics simulations, implied timescale plot and macrostate definition generated by PCCA+ clustering, time-trace of slow features, convergence plots for metadynamics simulations, free energy surface highlighting alternate Trp41 conformation, time-trace of different CVs for RIPK2 and RIPK2-XIAP complexes.

## Acknowledgements

The computations were performed on computer resources provided by the Swedish National Infrastructure for Computing (SNIC) at LUNARC (Lund University) and HPC2N (Umeå University).

