## Supporting Information for "AlphaFold-SFA: accelerated sampling of *cryptic pocket* opening, *protein-ligand* binding and *allostery* by AlphaFold, slow feature analysis and metadynamics"

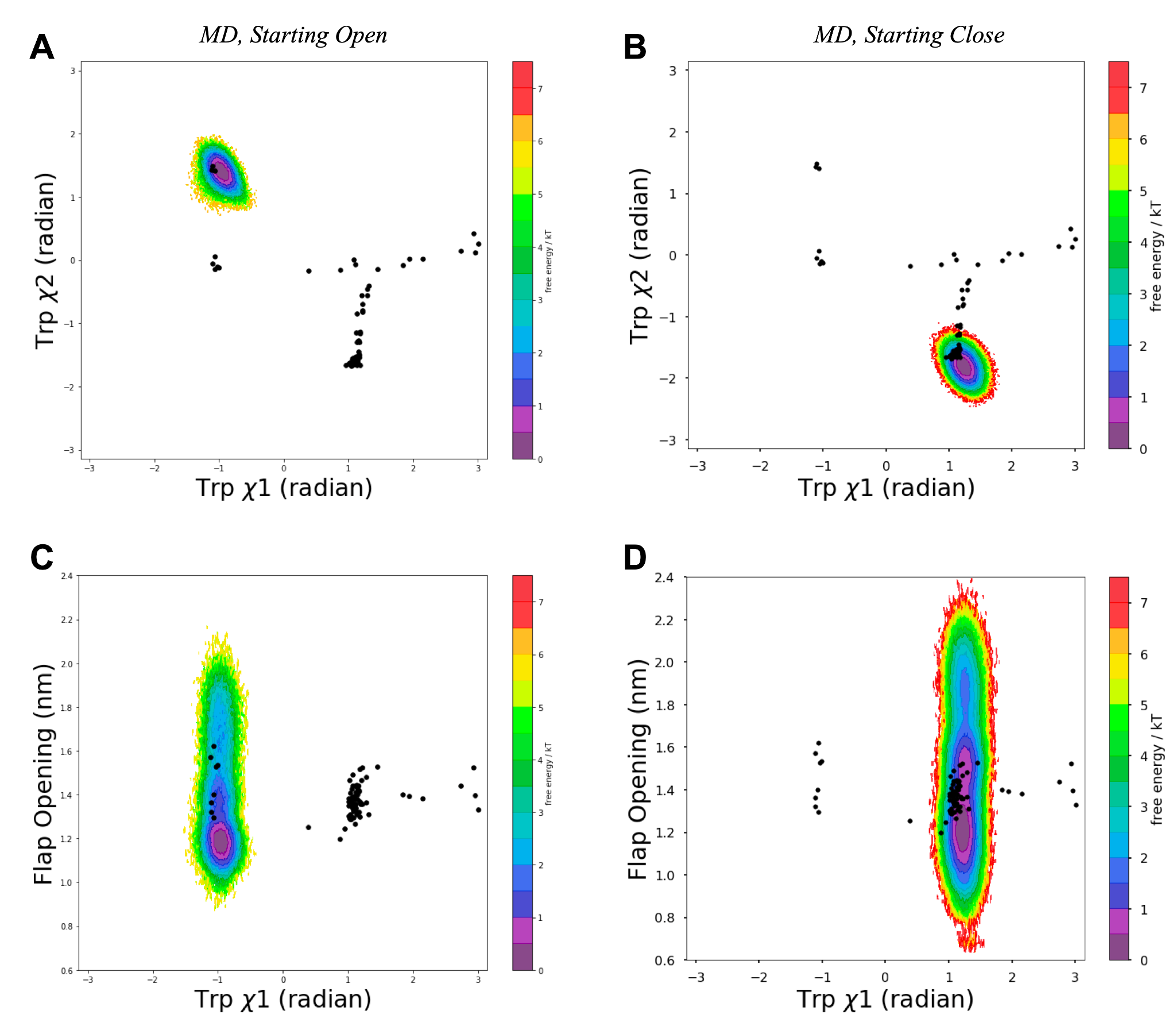

Figure S1. Unbiased molecular dynamics simulations with an aggregate simulation time of ~16 μs (160 replicas * 100 ns each) for apo-like open, PDB: 2BJU (A, C) and close, PDB: 1LF4 (B, D) plasmepsin II respectively, failed to sample multiple recrossing associated with Trp41 𝜒1 flipping. Projection of Trp41 𝜒1 angle along 𝜒2 and flap opening distance highlights the conformational heterogeneity captured in unbiased molecular dynamics simulations.

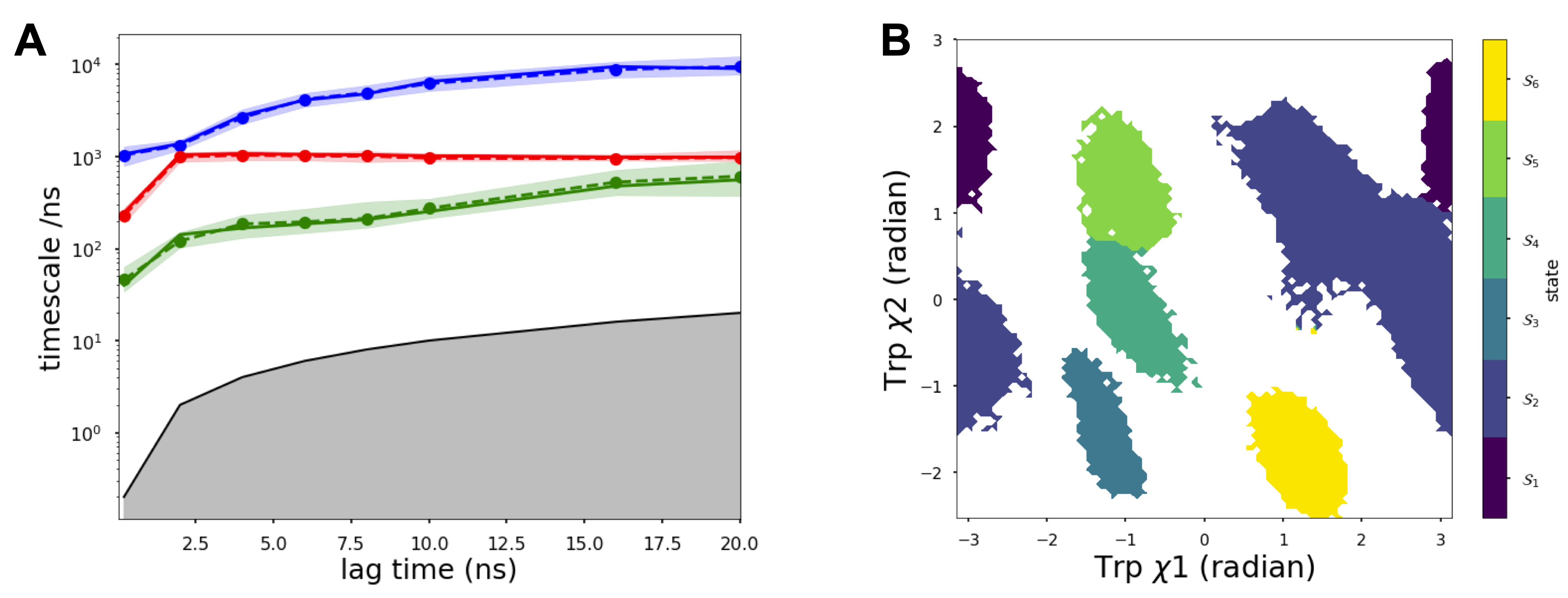

Figure S2. Implied timescale plot associated with Markov state model (A). We chose lag time of 6ns to generate MSM. *PCCA^+^* was used to generate microstate definition associated with Trp41 𝜒1 and 𝜒2 angles (B). *PCCA^+^* manage to separate close (*S6*) and open (*S5*) states in plasmepsin II.

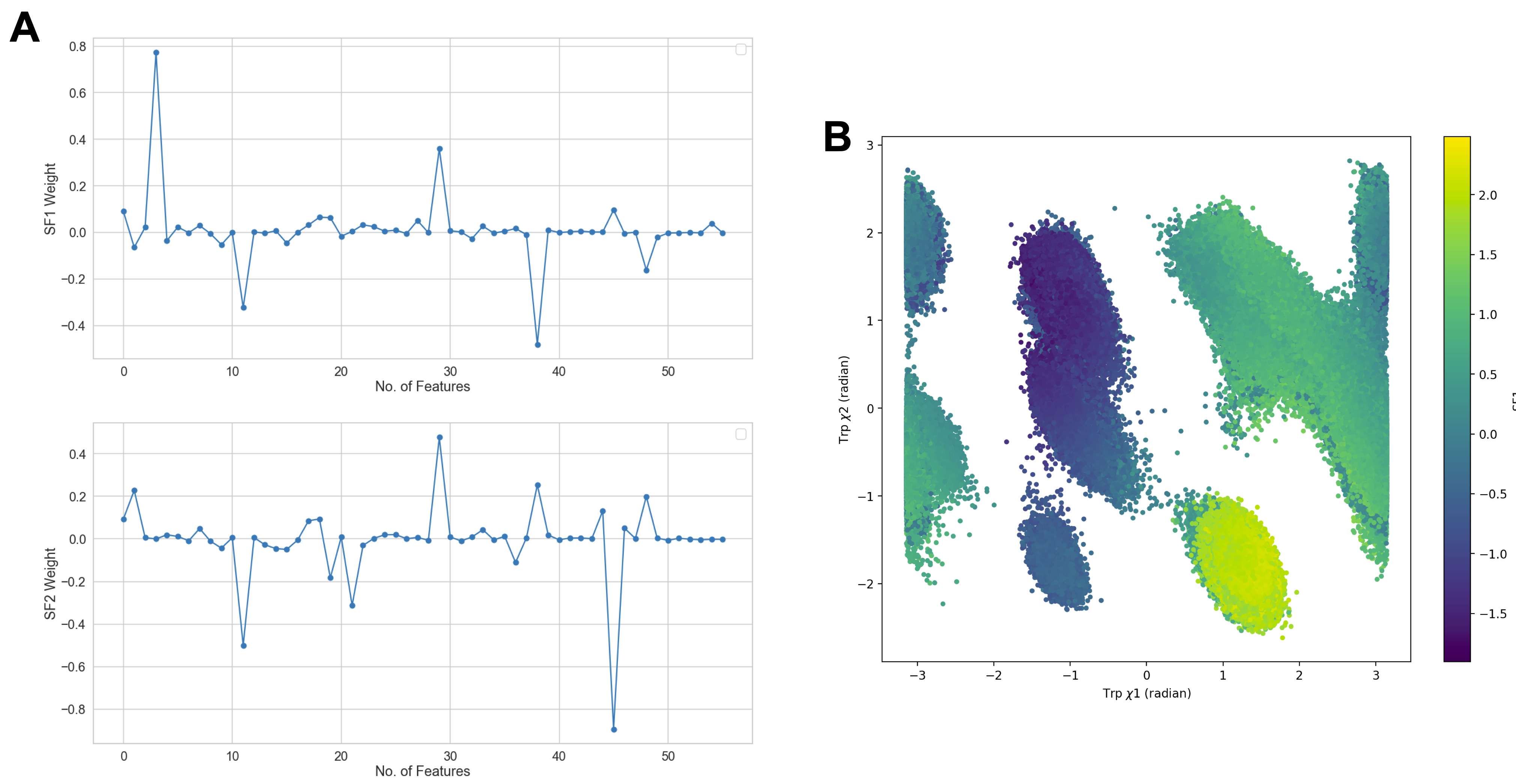

Figure S3. Weights corresponding each features showed how first two slow features (A) manage to capture sidechain flipping associated with Trp41 and Tyr77, 𝜒1 and 𝜒2 angles (see XXX for list of features and corresponding weights). Projection of Trp41 𝜒1 and 𝜒2 angles along SF1 highlights how SF1 manages to separate close and open states in plasmepsin II (B).

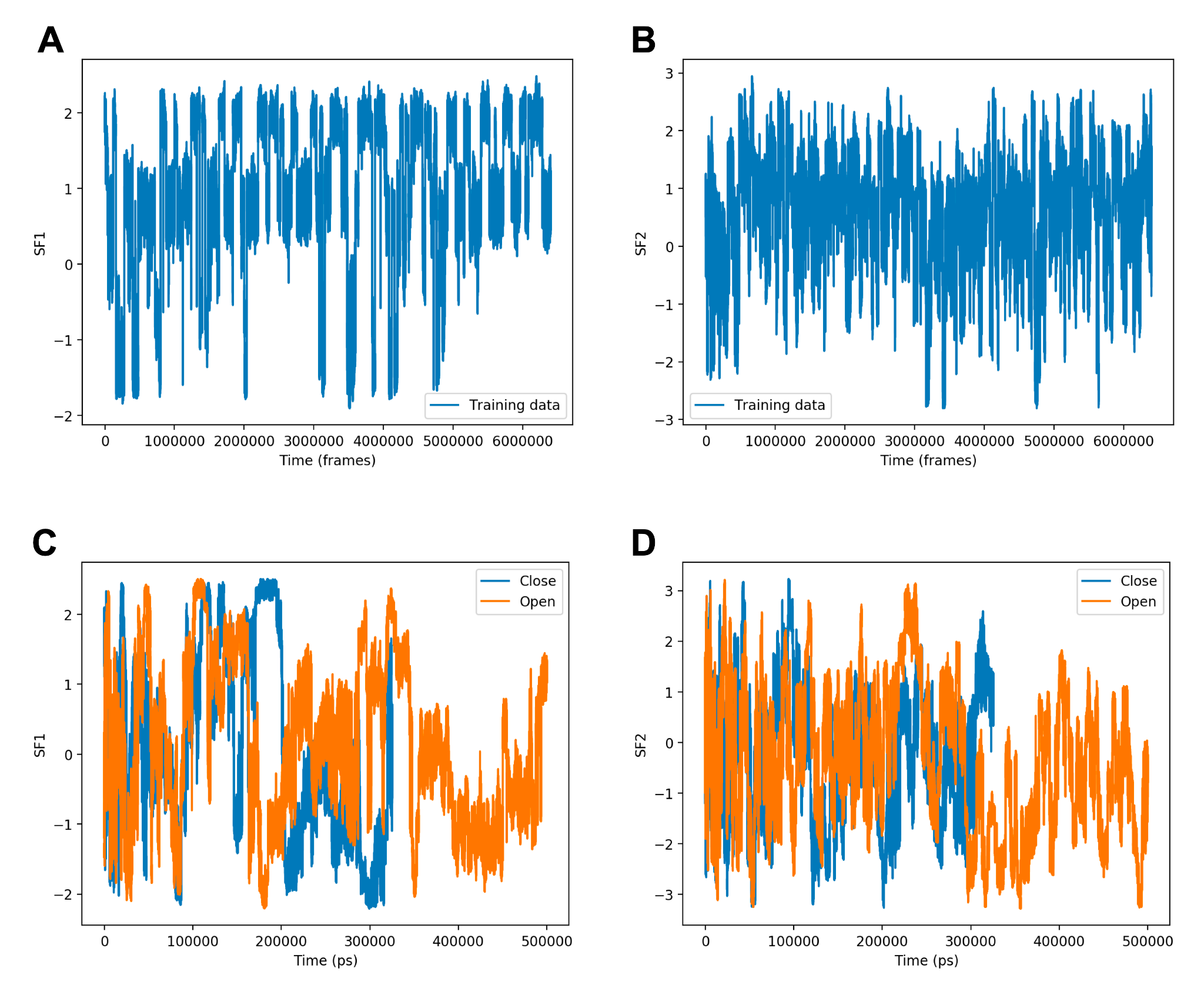

Figure S4. First two slow features projected on the training data (A, B) generated by unbiased molecular dynamics simulations launched from AlphaFold generated ensemble. Metadynamics simulations using slow features as CVs manage to capture multiple recrossing within a few hundreds of nanoseconds (C, D).

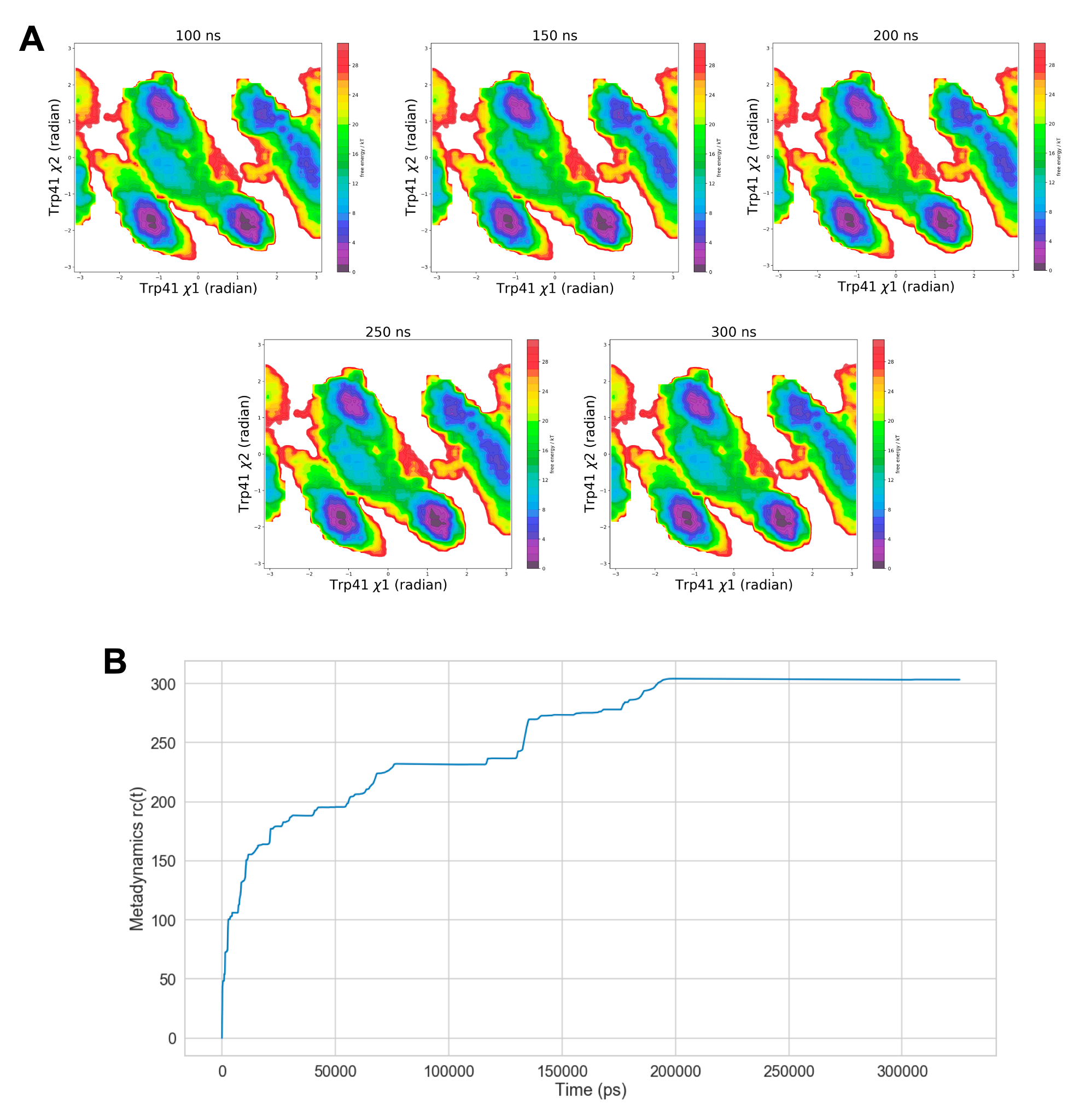

Figure S5. Reweighted free energy surfaces from well-tempered metadynamics projected along Trp41 𝜒1 and 𝜒2 angles at different time intervals highlighted convergence of our metadynamics simulations for the choice of force field and water model (A). Time trace of the reweighting factor *rc(t)* also highlighted the convergence of metadynamics.

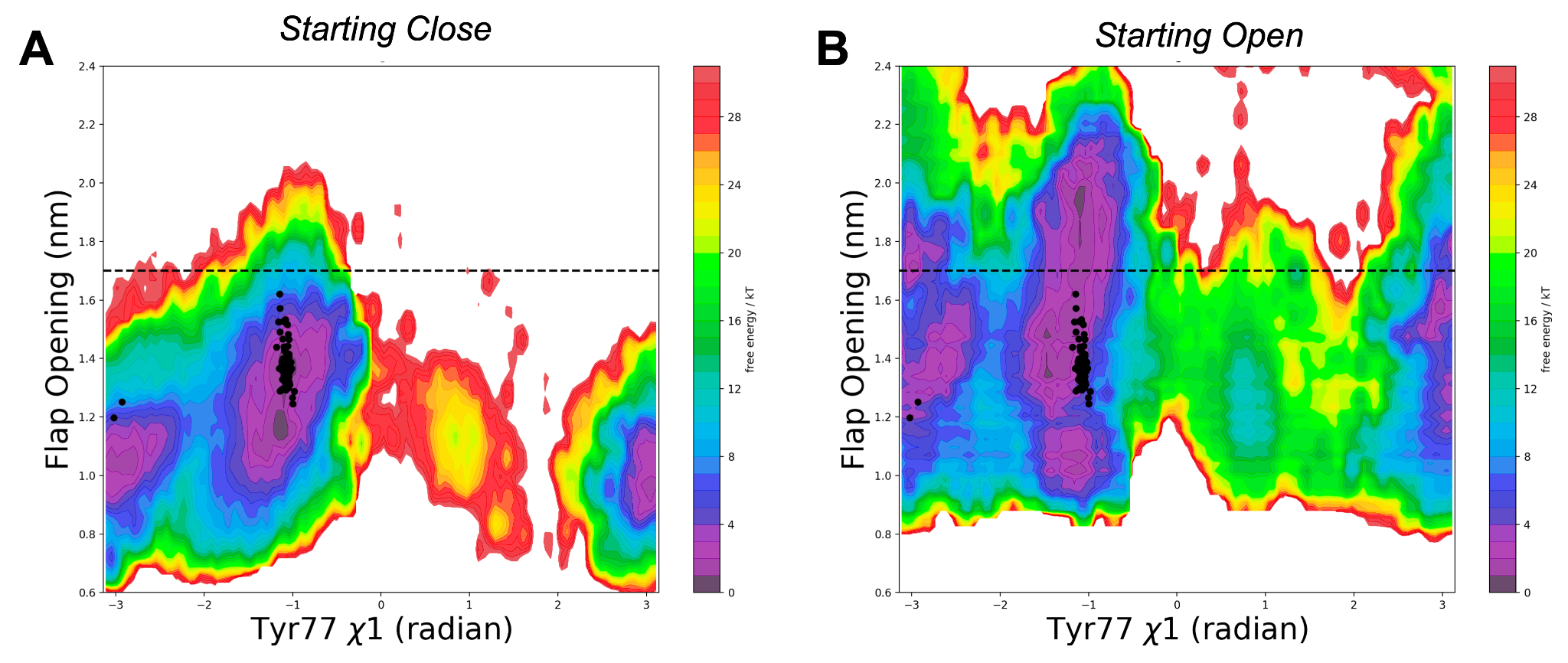

Figure S6. Reweighted free energy surfaces from SFA-metadynamics highlight how flipping of Tyr77 𝜒1 angles governs flap opening as discussed by Bhakat & Söderhjelm. AlphaFold generated conformations are shown as black dots.

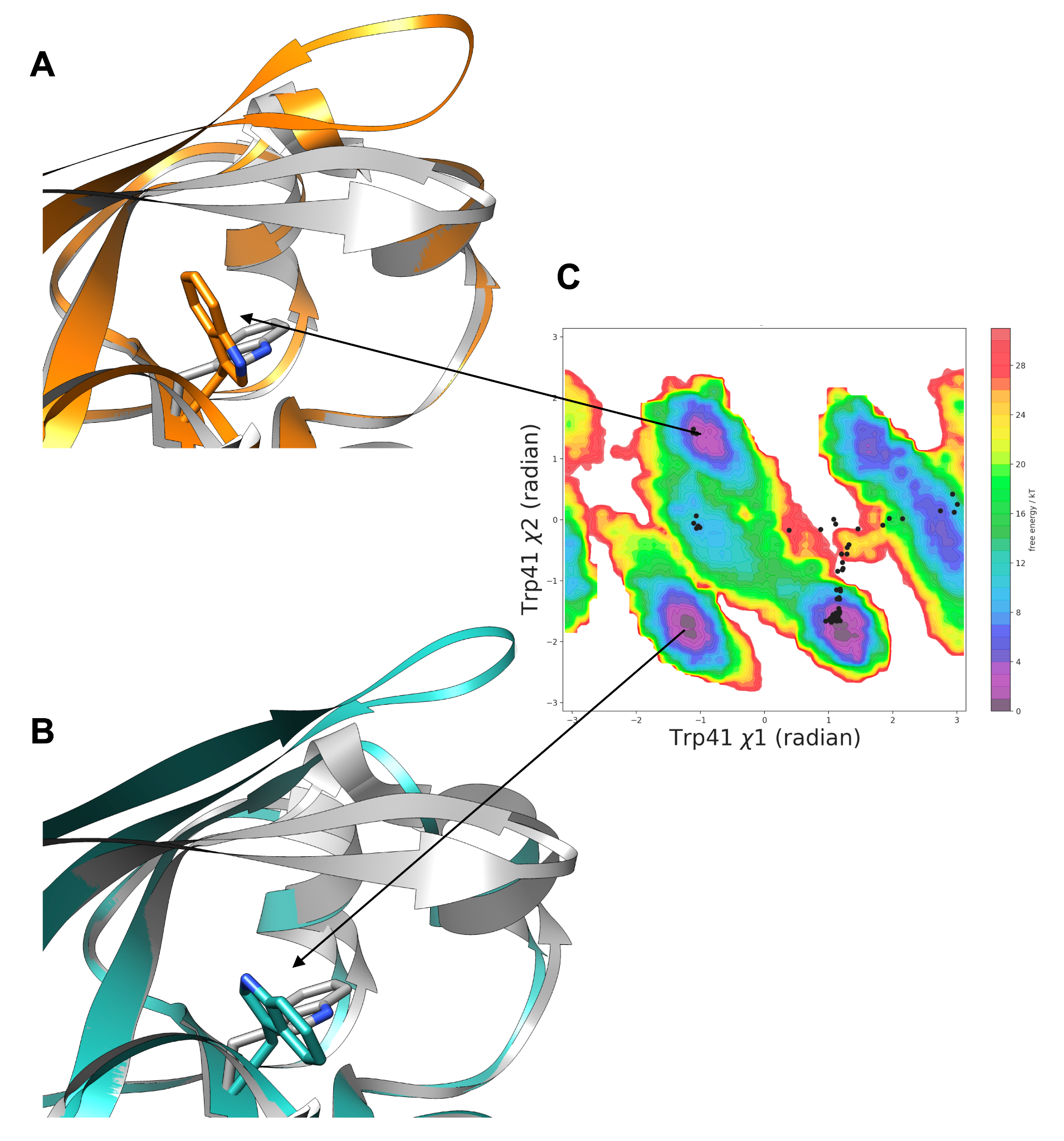

Figure S7. Reweighted free energy surface from SFA-metadynamics (C) along Trp41 𝜒1 and 𝜒2 angles highlighted alternate states (A, B) associated with 𝜒2 flipping. Orange (PDB: 2BJU) and blue (PDB: 4Z22) corresponds to deep cryptic pocket open states with 𝜒2 angle of +1 and -1 radian accordingly. AlphaFold ensembles are highlighted in black dots. It is important to note that AlphaFold failed to sample an alternate Trp41 𝜒2 angle of -1 radian.

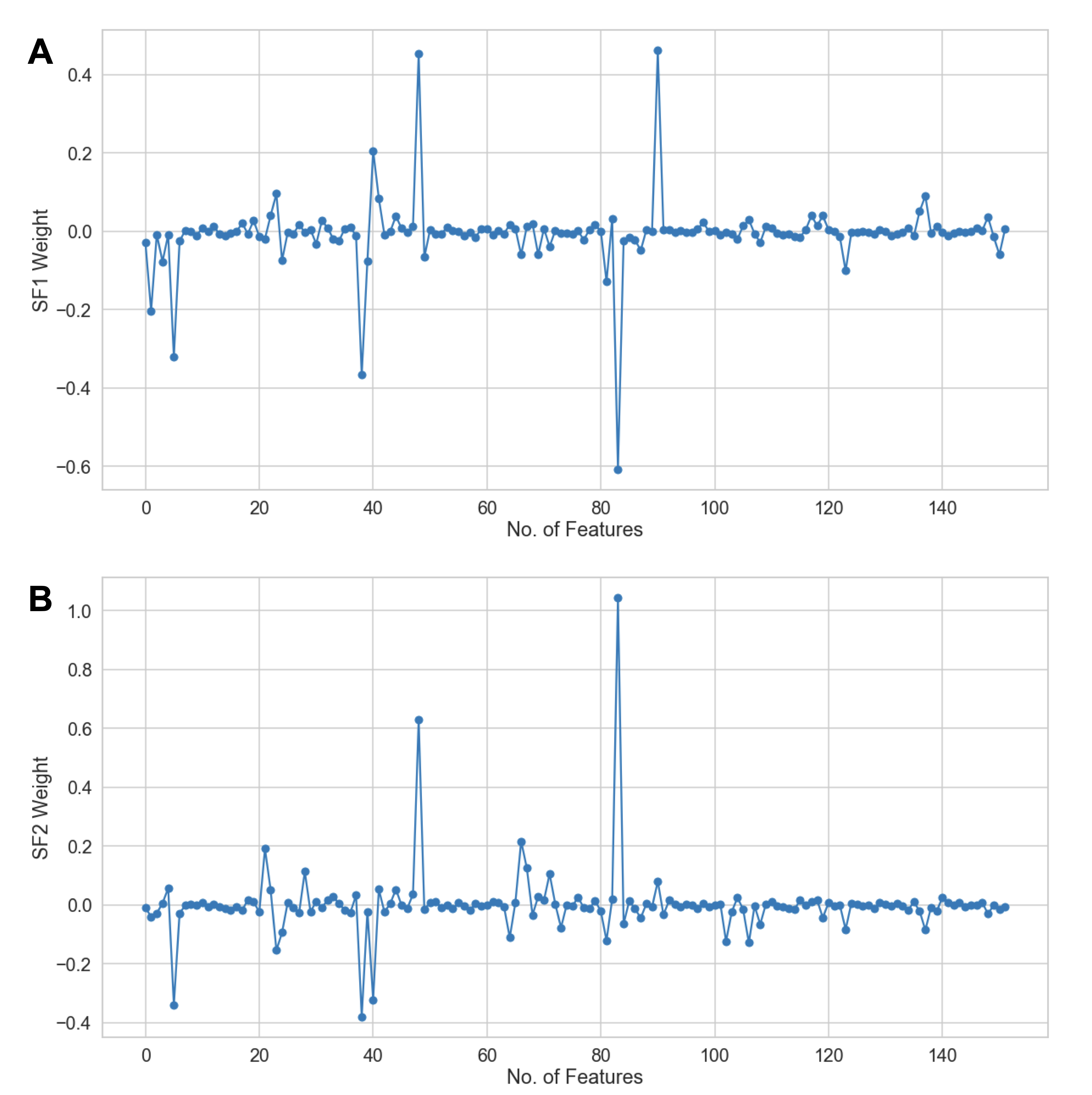

Figure S8. Weights corresponding first two slow features for sin/cos transformed dihedral angles associated with amino acid residues in RIPK2.

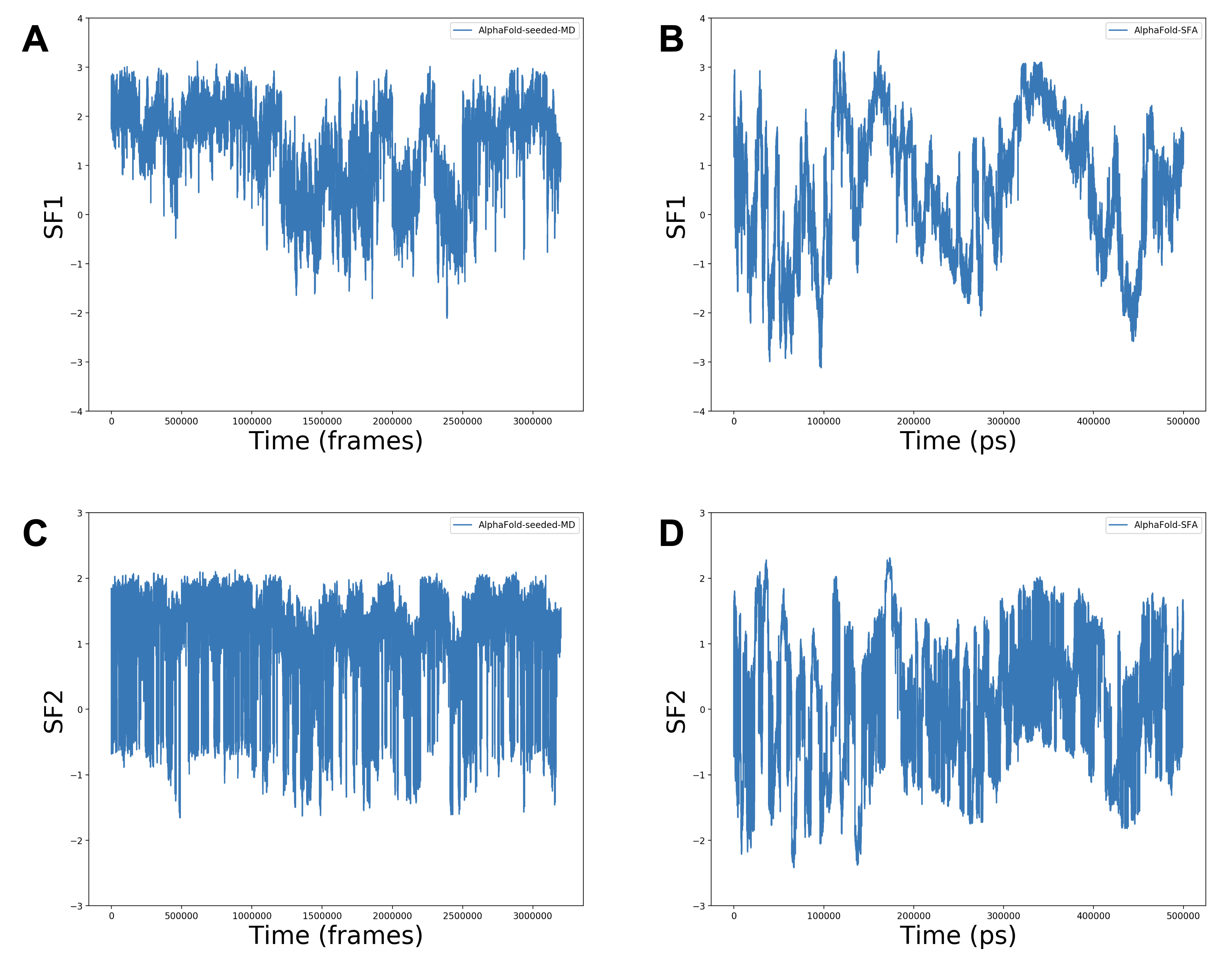

Figure S9. Sampling of first two slow features in AlphaFold-seeded MD and SFA-metadynamics simulations. Metadynamics accelerated the sampling along first two slow features which enabled sampling of allosteric dynamics in RIPK2.

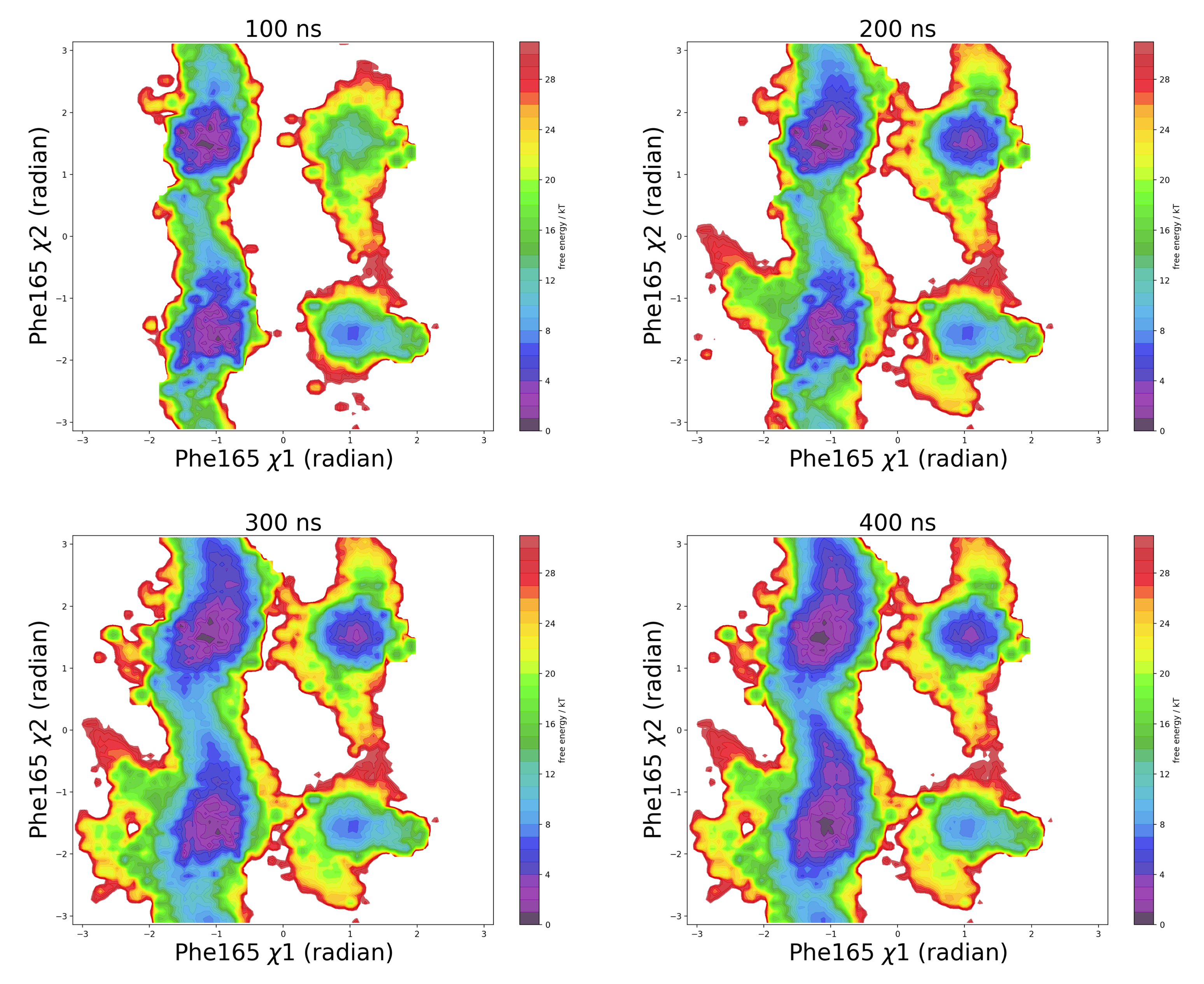

Figure S10. Reweighted free energy surfaces from well-tempered SFA-metadynamics projected along Phe165 𝜒1 and 𝜒2 angles at different time intervals highlighted convergence of the simulation.

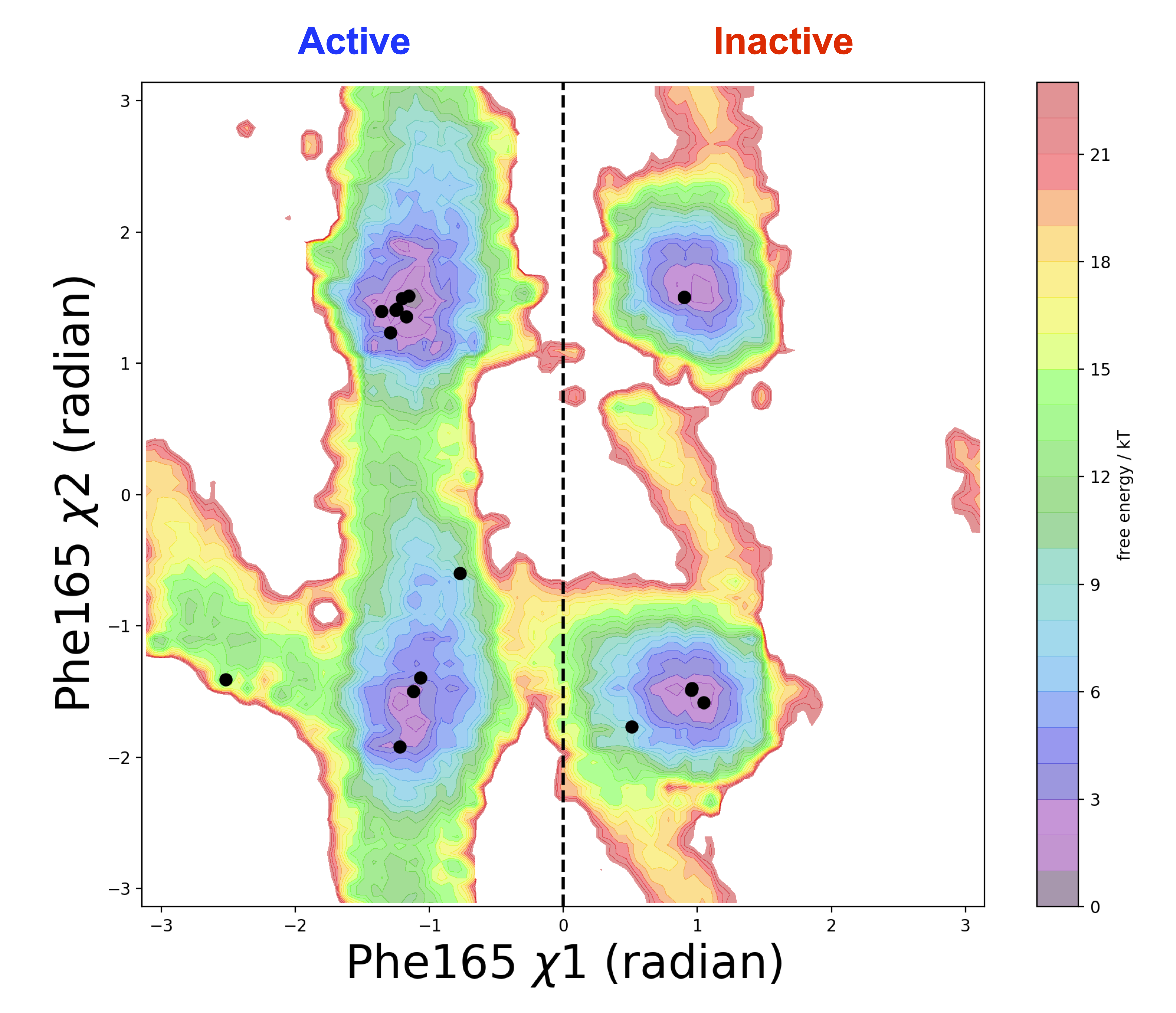

Figure S11. Distribution of Phe165 dihedral angles in RIPK2 crystal structures available in RCSB PDB projected on reweighted free energy surface from SFA-metadynamics. It is important to highlight that in inactive RIPK2 (PDB: 5J7B, 5AR4, 6SZJ, 6FU5) the position of Trp170 is not resolved.

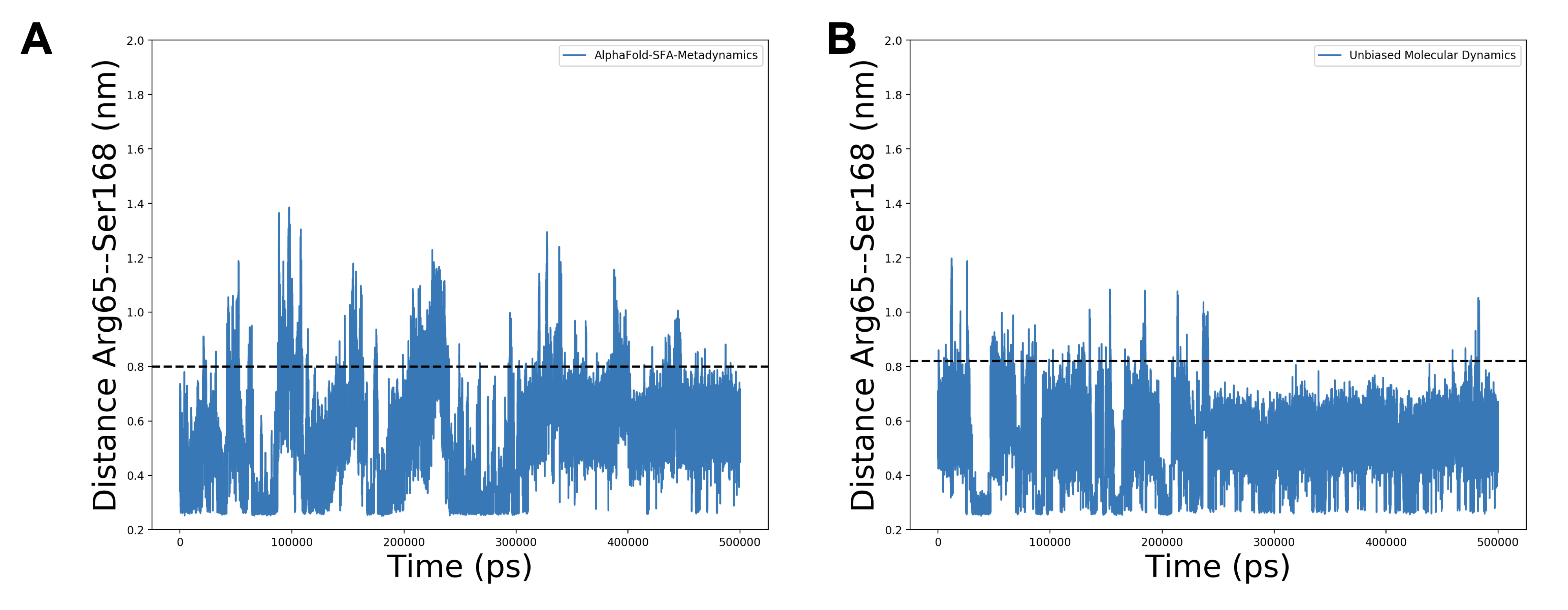

Figure S12. Time trace of Arg65—Ser168 distance during SFA-metadynamics (A) and unbiased MD simulation of apo RIPK2 (B). SFA-metadynamics managed to sample multiple transitions between active and inactive states of RIPK2 (demarked by the dashed line at 0.82 nm) compared to unbiased MD simulation which remained in the active state.

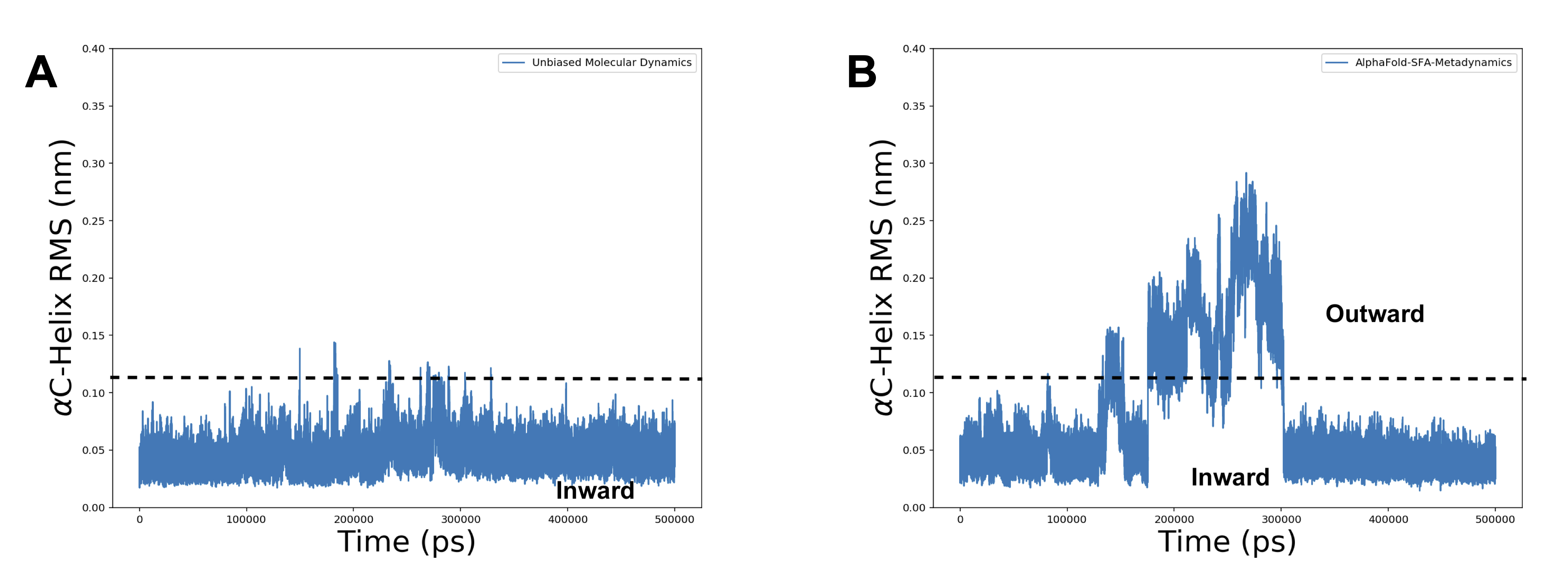

Figure S13. RMSD of αC helix during unbiased MD (**A**) and SFA-metadynamics (**B**) simulations. In unbiased MD simulations αC helix remained in an *inward* conformation due to H-bond interaction involving Arg65—Ser168. SFA-metadynamics simulations managed to capture conformational dynamics associated with the activation loop of RIPK2 which sampled flipping of Trp170. Flipping of Trp170 destabilizes the activation loop which breaks Arg65—Ser168 interaction and resulted in an *outward* conformation of αC helix.

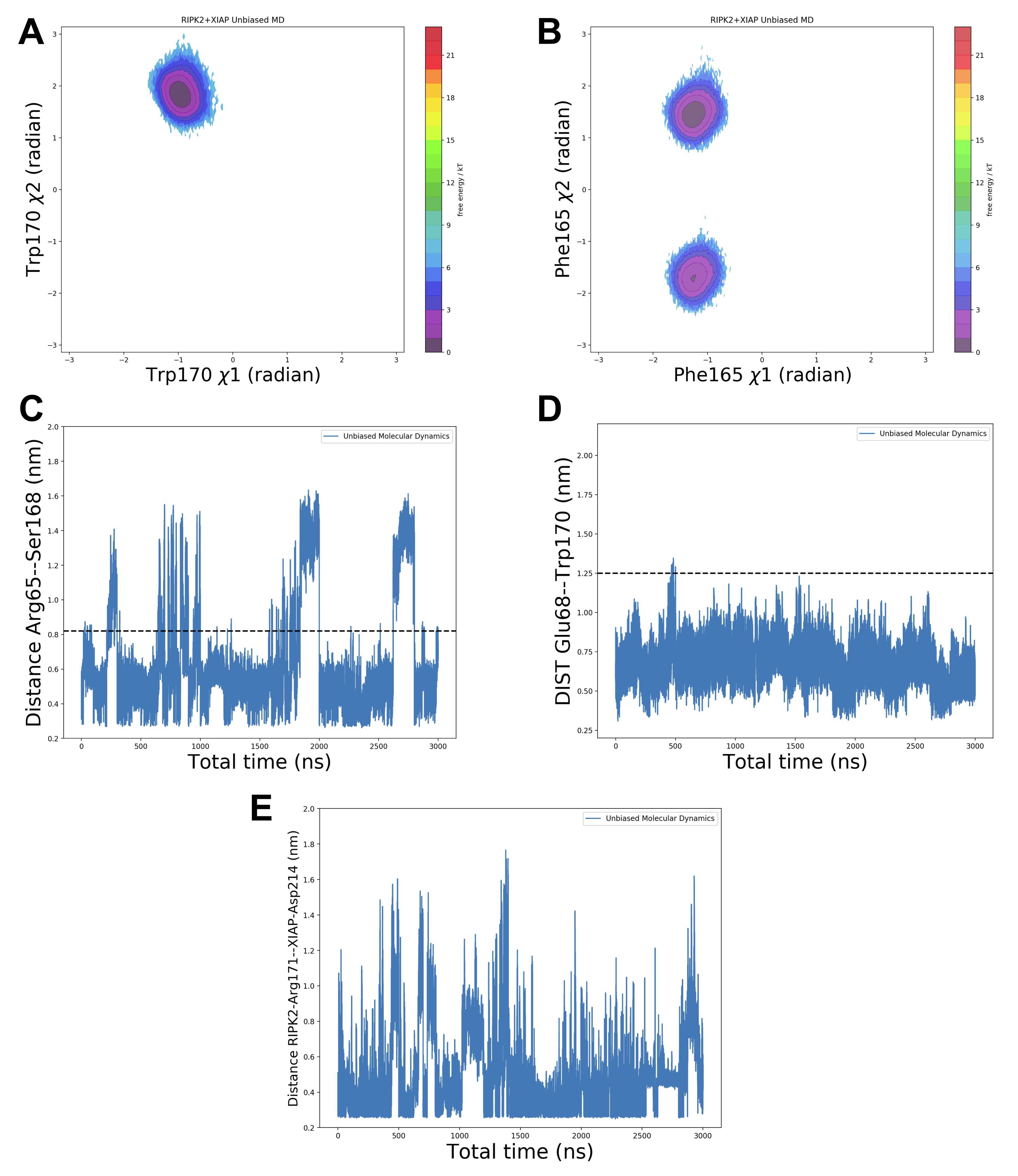

Figure S14. Population distribution of Trp170, Phe165 dihedral angles and distances between Arg65—Ser168, Glu68—Trp170 and Arg171—XIAP-Asp214 during total 3 μs of unbiased molecular dynamics simulations started from the cryo-EM model of XIAP-RIPK2. Simulations highlight that RIPK2 remains in an active state during RIPK2-XIAP interaction.

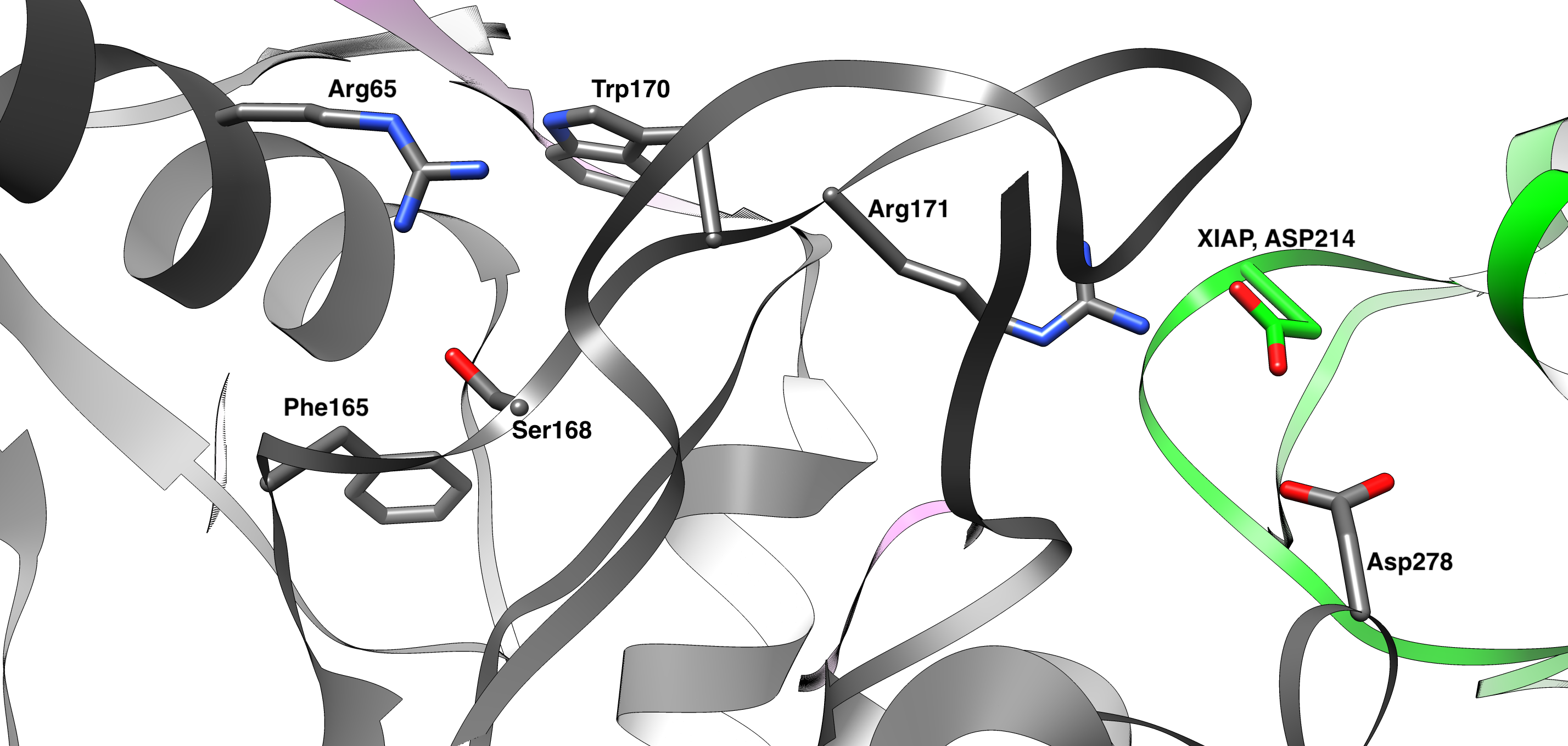

Figure S15. Structural insights of how Arg171 present in the activation loop of RIPK2 interacts with the Asp214 of XIAP (green). It also highlights orientation of Phe165, Trp170, Arg65 and Ser168, key residues involved in conformational dynamics of RIPK2.

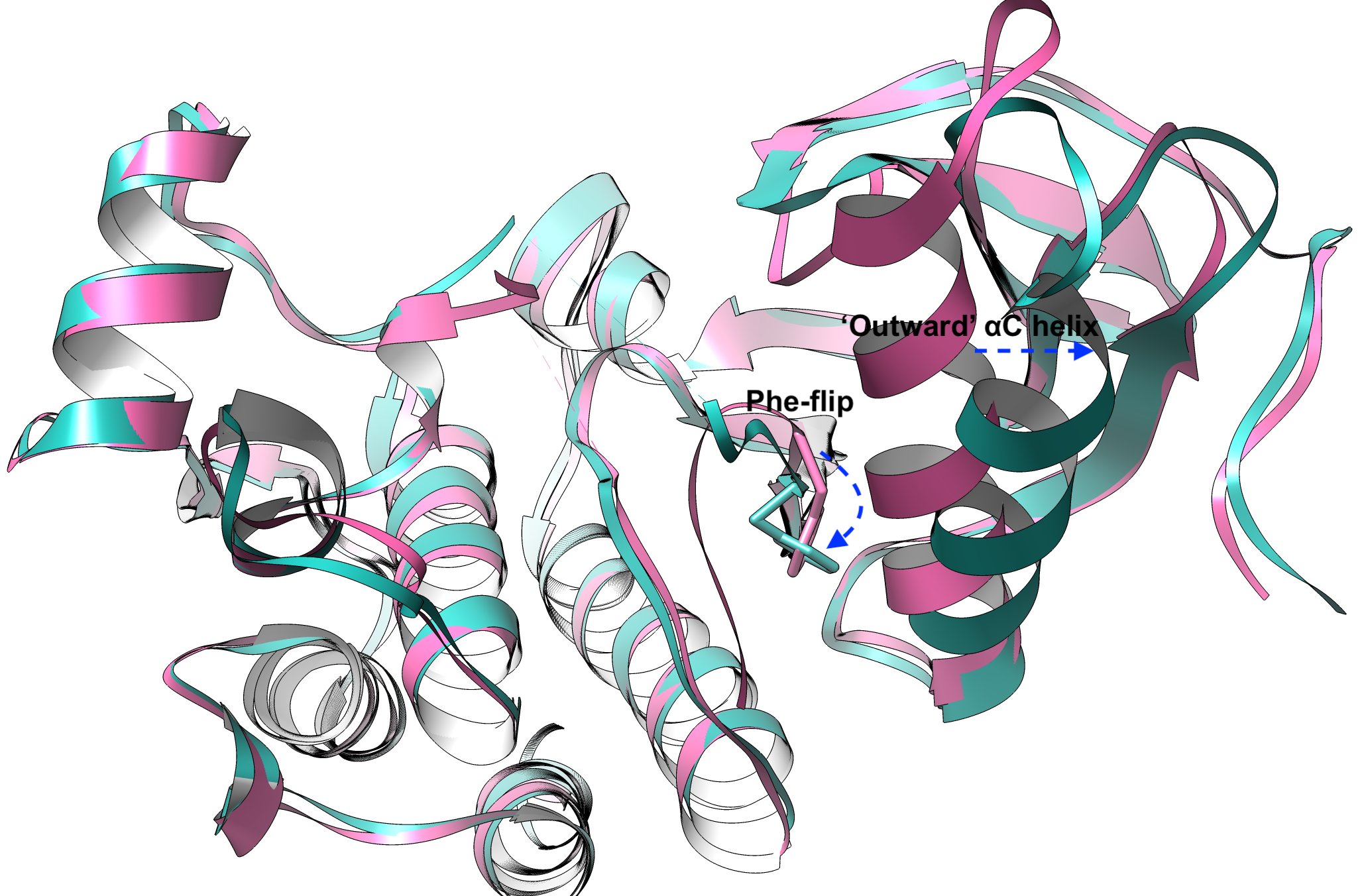

Figure S16. Crystal structures of holo-BRAF highlights the coupled motion involving DFG-Phe flipping and the ‘outward’ conformation of αC helix. Blue indicates PDB: 4EHG and the magenta indicates PDB: 2FB8.

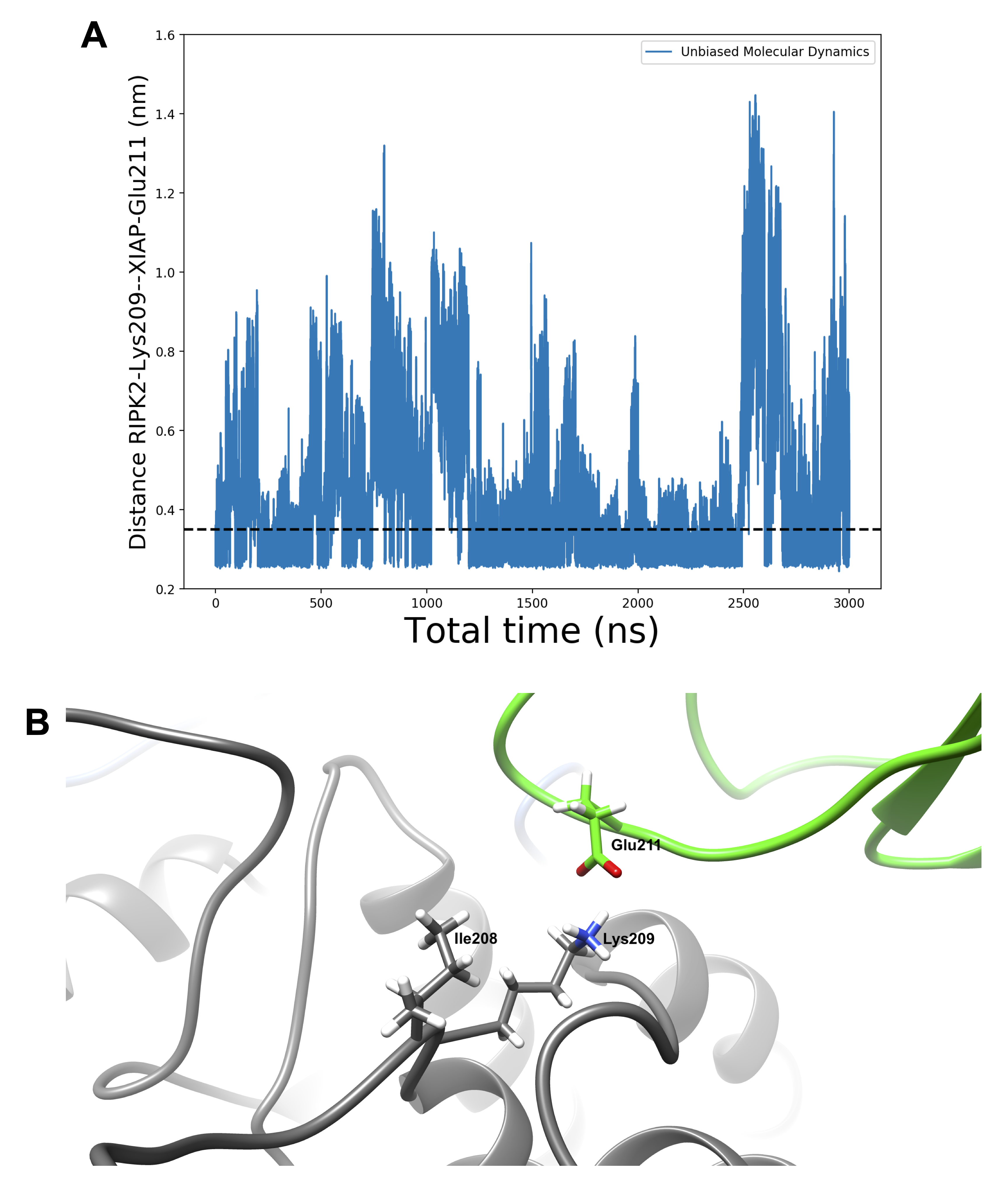

Figure S17. Time-trace of distance between Lys209 of RIPK2 and Glu211 of XIAP during total 3 μs of unbiased MD simulation of XIAP-RIPK2 complex (A). The dashed line at 0.35 nm indicates formation of H-bond interaction involving Lys209—Glu211. Orientation of Lys209 and Ile208 of RIPK2 and Glu211 of XIAP is highlighted for visual inspection (B).

**RIPK2-XIAP interface**

Molecular dynamics simulation of RIPK2-XIAP complex highlighted key residues in the protein-protein interface stabilizing the complex (Figure S18 and Figure S19).

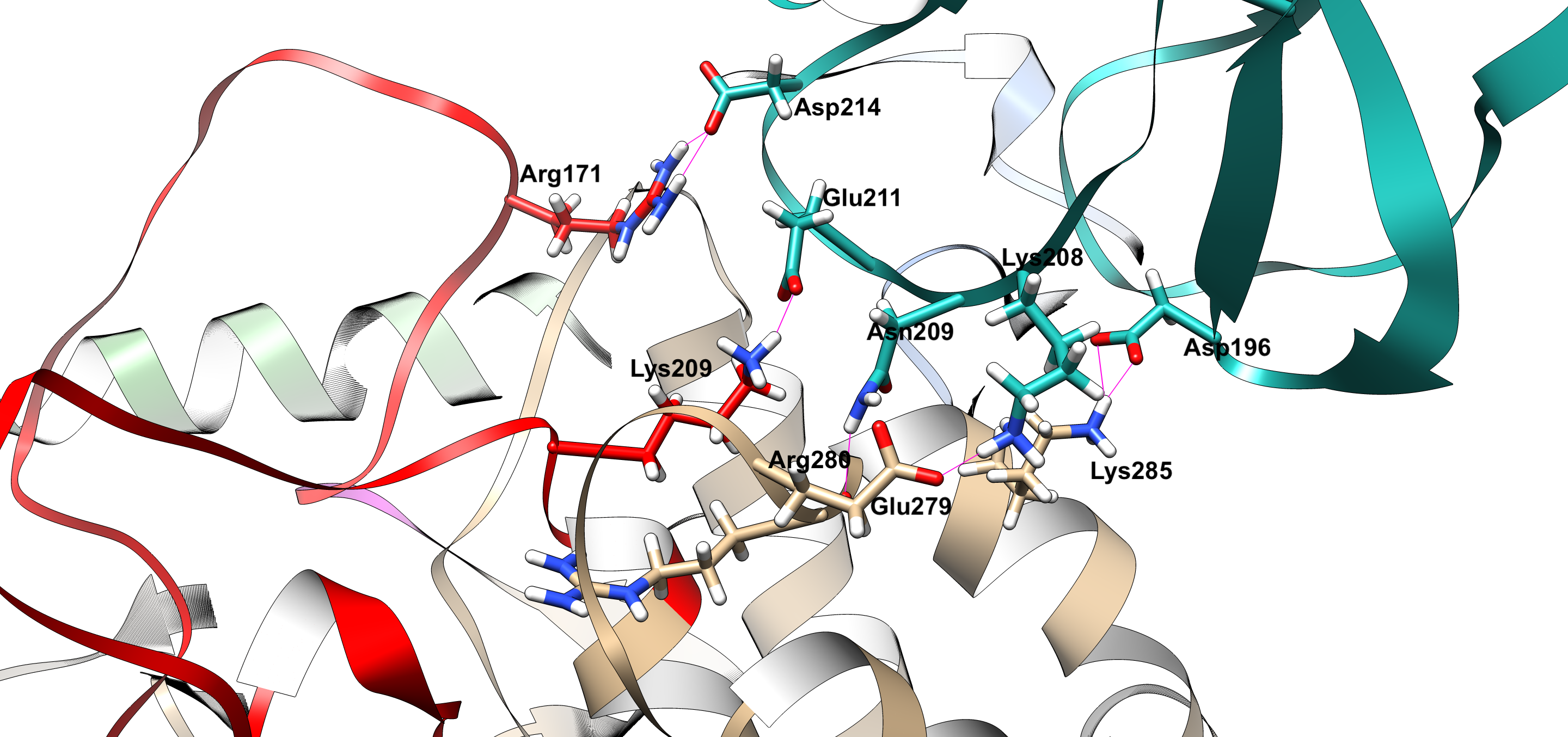

Figure S18. PPI interface of RIPK2-XIAP complex highlighting key residues involved in H-bond interactions. The activation loop of the RIPK2 is highlighted in red and the XIAP is highlighted in ‘seagreen blue’.

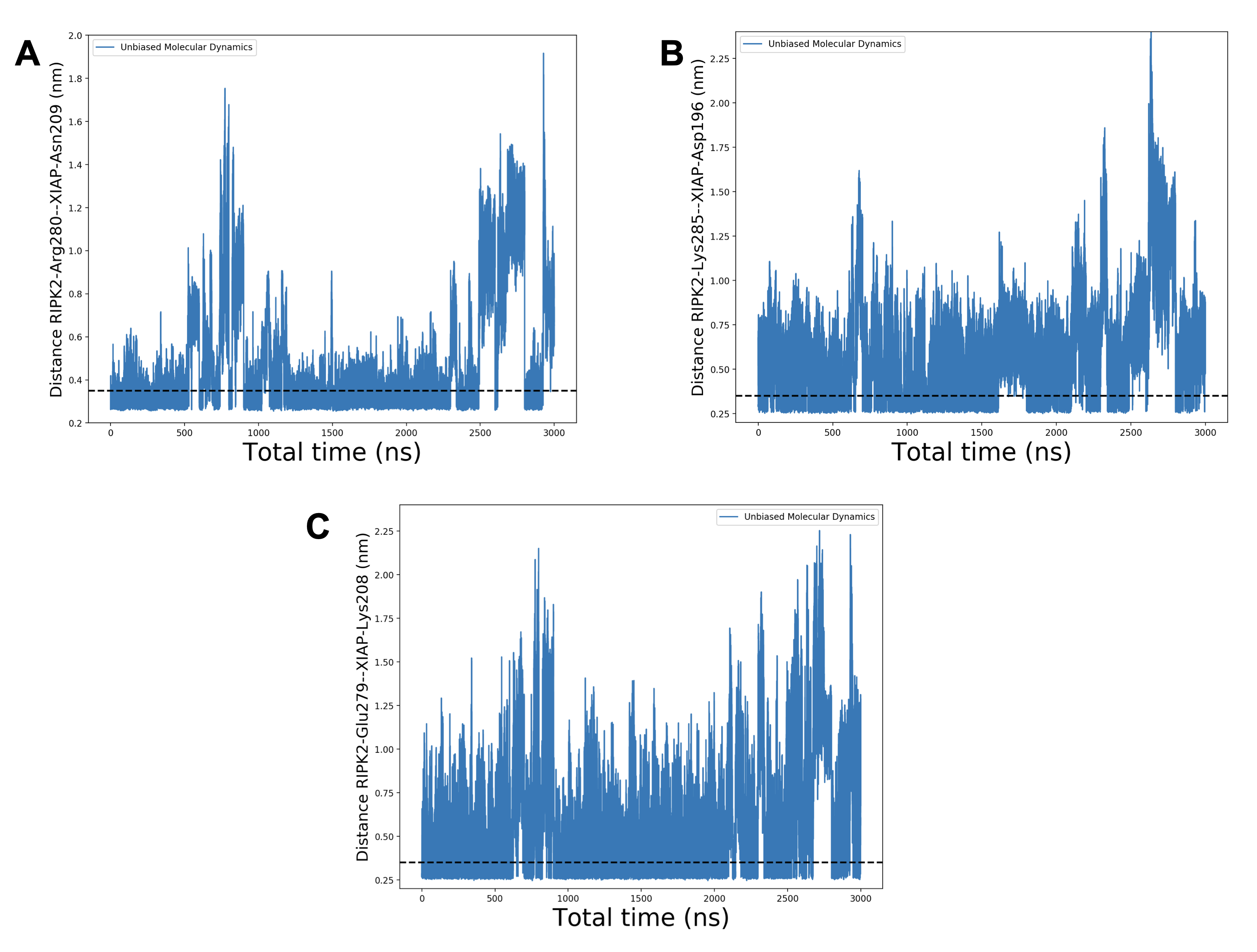

Figure S19. Temporal evolution of hydrogen bond distances throughout molecular dynamics simulations, encompassing 15 independent runs each lasting 200 nanoseconds, across crucial residues that play a role in stabilizing the RIPK2-XIAP complex. The dotted line at 0.35 nm indicates complete formation of H-bond.
